# Indian Red Jungle fowl depicts close genetic relationship with Indian native chicken breeds as evidenced through whole mitochondrial genome intersection

**DOI:** 10.1101/2020.12.29.424655

**Authors:** M. Kanakachari, R.N. Chatterjee, U. Rajkumar, S. Haunshi, M. R. Reddy, T.K. Bhattacharya

**Affiliations:** ICAR-Directorate of Poultry Research, Rajendranagar, Hyderabad, India

**Keywords:** Chicken, Mitochondrial DNA, Next-generation sequencing, SNPs, Mutations and variants, Molecular phylogeny

## Abstract

Native chickens are dispersed in a wide range of geometry and they have influenced hereditary assets that are kept by farmers for various purposes. Mitochondrial DNA (mtDNA) is a widely utilized marker in molecular study because of its quick advancement, matrilineal legacy, and simple molecular structure. In a genomics study, it is important for understanding the origins, history, and adjustment of domestication. In this report, for the first time, we utilized Next-generation sequencing (NGS) to investigate the mitochondrial genomes and to evaluate the hereditary connections, diversity, and measure of gene stream estimation in seven Indian native chicken breeds along with twenty-two Asian native breeds. The absolute length of each mtDNA was 16775bp harboring 4 tRNA genes, 2 rRNA genes, 12 protein-coding genes, and 1 D-loop region. The chicken breeds were genotyped by using the D-loop region and 23 haplotypes were identified. In addition, when compared to only Indian native breeds more haplotypes were identified in the NADH dehydrogenase subunit (ND4 and ND5), Cytochrome c oxidase subunit (COXI and COXII), Cytochrome b, mitochondrial encoded ATP synthase membrane subunit 6, and Ribosomal RNA genes. The phylogenetic examination utilizing N-J computational algorithms indicated that the analyzed all native chicken breeds were divided into six significant clades: A, B, C, D, E, and F. All Indian native breeds are coming under the F clade and it says all Indian breeds are domesticated in India. Besides, the sequencing results effectively distinguished SNPs, INDELs, mutations, and variants in seven Indian native breeds. Additionally, our work affirmed that Indian Red Jungle Fowl is the origin of reference Red Jungle Fowl as well as all Indian breeds, which is reflected in the dendrogram as well as network analysis based on whole mtDNA and D-loop region. Albeit, Indian Red Jungle Fowl is distributed as an outgroup, proposing that this ancestry was reciprocally monophyletic. The seven Indian native chickens of entire mtDNA sequencing and disclosure of variations gave novel insights about adaptation mechanisms and the significance of important mtDNA variations in understanding the maternal lineages of native chickens.

## 1. Introduction

Modern day’s chicken breeds are mostly evolved from Red Jungle fowl, which is also evident from archeological discoveries [1–5]. There are also some reports of the contribution of other Jungle fowl in evolving many breeds across the Globe [6,7]. However, chicken domestication has first occurred, the evidence for times and places are disputable [8–14]. In early 3,200 BC, the chicken domestication was observed in the Indus valley and accepted the epicenter of chicken domestication [8]. Afterward, brought up issues about the exclusive domestication at Indus valley after excavations at Peiligan Neolithic sites in China and proposed the alternate and conceivable prior domestication centers [10]. It is recommended that the wild RJF found in the forests of South East Asia and India, when people domesticated the chicken and spread to different parts of the world, the genome landscape of domestic chicken was molded by natural and artificial selections, bringing about a wide range of breeds and ecotypes [12,15–19]. The domestic chicken has broad phenotypic variations yet missing in the Red Jungle Fowl are the result of domestication, for example, plumage color and other morphological characteristics, behavioral, and production traits, adaptation to different agro-ecosystems and rigid human selection for production as well as aesthetic qualities [20–22].

Following domestication, the enormous breeding programs have come about more than sixty chicken breeds representing four particular genealogies: egg-type, game, meat-type, and bantam [23]. Whereas, some authors propose a monophyletic origin of domestic chicken and others give proof for multiple and independent domestication events [5,11,12]. According to the taxonomy, the genus Gallus is composed of four species, they are G. gallus (Red Jungle Fowl), G. lafayettei (Lafayette’s Jungle Fowl), G. varius (Green Jungle Fowl), and G. sonneratii (Grey Jungle Fowl). At Present, Red Jungle Fowl has five sub-species based on phenotypic traits and geographic distribution of the populations, such as G. g. gallus (South East Asia Red Jungle Fowl), and G. g. spadiceus, G. g. bankiva, G. g. murghi (Indian Red Jungle Fowl) and G. g. jabouillei [24]. In publications, wild and domesticated birds are frequently referred to as ‘fowls’ and ‘chicken’, respectively. The domestic chicken is considered either as a subspecies of RJF (G. g. domesticus) or as a separate species, G. domesticus. However, tight clustering of the distinctive subspecies limits this current taxonomical chain rendering sub-species status within RJF redundant [11].

Other than the taxonomical details, the scientists are likewise worried about the RJF genetic integrity and conservation status of the wild and those held in avicultural assortments. It was revealed that the domestic chicken is hybridized with the wild RJF resulting in erosion of the genetic purity of the wild birds [16,25,26]. Because the previous examinations depend on phenotypic characters, or small samples are utilized for DNA investigations. Mitochondrial D-loop sequence phylogeny and nuclear gene analyses demonstrated conceivable hybridization between GJF-RJF/domestic birds [26]. Based on these reports, the evaluation of the genetic uniqueness of Indian RJFs is significant for conservation and population studies.

Chickens are widespread fowls and have been domesticated in India and South-East Asia for various purposes including decoration, religions, cockfighting, and food production [22,27]. In India, village farmers grow chickens for food, so, according to the local environment circumstance their morphology, behavior, physiology, and adaptation have been changed [28]. India is a huge nation that contains a unique scope of altitudes and climates, because, native chickens have an astounding genetic diversity. In the twentieth century, the dispersion of chicken genetic assets resources in general inimically affected the industrialization and globalization of chicken production. In this way, various chicken varieties have recently been vanished or really at risk of elimination. As indicated by ICAR-NBAGR, India has 19 indigenous breeds, they are Ankaleshwar, Aseel, Busra, Chittagong (Malay), Danki, Daothigir, Ghagus, Harringhata Black, Kadaknath, Kalasthi, Kashmir Favorolla, Miri, Nicobari, Punjab Brown, Tellichery, Mewari, Kaunayen chicken, Hansli and Uttara. A huge number of streams of various sizes, shapes, and hues, and generally taking after the wild fowls, are found all over India. They change in appearance as indicated by the region in which they have been reproduced. Out of the 19 enrolled native chicken types of India, eight unique lines/breeds used in the current examination, they are Aseel, Ghagus, Nicobari (brown and black), Kadaknath, Haringhata black, Red Jungle fowl, and Tellicherry. There is a need to characterize native chicken lines at the molecular level to make protection and improvement activities to benefit the nation’s people. With the extended focus on genetic preservation, remarkable alleles may be helpful in choices to keep up native varieties. The information on genetic decent variety examinations of the chicken germplasm being used for the advancement of rural chicken varieties is small. Thus, the examination was endeavoring to choose the genetic heterogeneity between chicken populaces, which are being used in the progression of provincial chicken assortments in India.

Mitochondria assume a significant role in aerobic respiration through oxidative phosphorylation producing most of cellular ATPs as definitive oxygen transduction, by spending 90% of cellular oxygen take-up [29,30]. To decide the role of mitochondrial genes in high-altitude adaptation, six high-altitude phasianidae birds and 16 low-altitude relatives of mitochondrial genomes were analyzed and revealed significant evidence for positive selection in the genes viz. ND2, ND4, ND5, and ATP6 [31,32]. For high-altitude adaptation, cytochrome c oxidase (COX) was the key mitochondrial gene and assume a significant role in oxidative phosphorylation regulation and oxygen sensing transfer, thus, it is thinking about that the MT-COI and MT-CO3 genes have an association with adaptation [33–35]. Analyzing the role of the ATP-6 gene in the proton translocation and energy metabolism, estimated the possibility of mutation assuming a significant role in easier energy conversion and metabolism to better adapt to the harsh environment of the high-altitude areas [35]. In addition to enhancing oxygen uptake and transport for adaptation to high-altitude hypoxic conditions, it is essential to improve the effectiveness of oxygen utilization. Subsequently, the mitochondrial genome, which encodes 13 basic oxidative phosphorylation system proteins (7 NADH dehydrogenase complex subunits, cytochrome bc 1 complex cytochrome b subunit, 3 cytochrome c oxidase subunits, and 2 ATP synthase subunits), must have experienced natural selection during adaptation to hypoxic conditions at high altitude [36].

Mitochondria assume a role in metabolism, apoptosis, disease, aging, signaling, metabolic homeostasis, macromolecules biosynthesis (lipids and heme), oxidative phosphorylation (for ATP synthesis) and other biochemical functions [37]. Mitochondria contain a class of cytoplasmic DNA molecules, i.e., mitochondrial DNA (mtDNA), it is maternally inherited, and there is no prompt evidence that it can recombine with other mtDNA molecules [26,38]. This suggests vertebrate mtDNA is going on through female lineages in a clonal way with no horizontal mixing and this makes it clearer to reproduce an evolutionary history than the nuclear genome [39]. The mtDNA is profoundly polymorphic (5-10 times quicker evolutionary rate) contrasted with the nuclear DNA most likely because of the absence of a replication repair system [40,41]. Every cell has hundreds to thousands of mtDNA duplicates and it has a very high mutation rate possibly need defensive histones, inefficient DNA repair systems, and constant exposure to mutagenic impacts of oxygen radicals. The chicken mitochondrial DNA is a circular molecule contains about 16.8 kb size like other vertebrate mitochondrial DNA (mtDNA), it encodes 13 proteins, 2 rRNAs, and 22 tRNAs [42].

The underlying assessment of chicken hereditary assets, the molecular marker techniques may give valuable data by estimation of the hereditary distance. In chicken hereditary diversity examines, microsatellites have been effectively utilized. For examining the hereditary connections between chicken populations, hereditary assorted diversity estimates utilizing the exceptionally polymorphic variable number of tandem repeat loci have yielded reliable and precise data. Over the previous decade, the utilization of maternally inherited mitochondrial DNA (mtDNA), particularly its complete displacement-loop (D loop) region, has expanded. To track genetic information about chicken ancestral breeds, demonstrating the phylogenetic relationship, genetic distance, and variability within and between populations, one of the most significant and amazing molecular tools, the nucleotide sequence of the mitochondrial D-loop region was used [43]. The mtDNA or an explicit part of mtDNA (e.g. D-loop) sequencing gives better precise data on evolution and hereditary diversity [5,44]. The D-loop region evolves much faster than different areas of the mtDNA and it does not encode a protein. More than 20 years, for phylogenetic examination mtDNA and especially D-loop sequences have been utilized [45]. Variation study was done utilizing the mtDNA D-loop region and HVI domain of 397 bp fragment for 398 African native chickens from 12 countries and discovered 12.59% polymorphic sites [46]. Likewise, 25 individuals from six native Chinese chicken populaces recorded variation rates to be 7.05% and 5.54%, respectively [24,47].

Right now, a generous mtDNA examination is dire to produce the baseline genetic information on matrilineal phylogeny, genetic diversity and distance, and variability of RJFs and native chickens in India. However, there is no previous report on Indian native chickens at the mitochondrial genomic level. Next-generation sequencing (NGS) has become a tool to depict genomic highlights [48,49]. Examining native chickens at the mitochondrial genomic level can explain the mitochondrial genomic premise of their disparities and the particular attributes of indigenous chickens can be accurately investigated. The comprehension of phylogeography will clarify the demographic history, origin, and extension of chicken species. To defeat the issue of parallel mutations and lineage exchange between different populaces, network analysis has enhanced phylogenetic trees [50]. Therefore, the current examination aims to assess the hereditary divergence between twenty-two Asian native breeds and seven native Indian chicken breeds (i.e. Aseel, Ghagus, Nicobari Brown, Tellicherry, Kadaknath, Haringhata Black and Red Jungle Fowl) utilizing mtDNA NGS sequence data.

## 2. Materials and methods

### 2.1. Experimental birds and sample collection

The seven Indian native chicken breeds namely, Aseel, Ghagus, Nicobari brown, Kadaknath, Tellicherry, Haringhata black, and Red Jungle fowl were studied for which blood samples of one female bird each of Aseel, Ghagus, Nicobari brown, and Kadaknath were collected from the experimental farms of Directorate of Poultry Research, Hyderabad while samples of Haringhata black and Red Jungle fowl were collected from the experimental farms of WBUAFS, West Bengal and CSKHPKVV, Palampur, respectively. The blood samples of Tellicherry were collected from the local farmers of Kerala state (Fig. 1; Table 1; Table 2). The blood samples were collected from adult birds of all the breeds and stored at −80°C. The DNA was extracted from blood samples according to the lab standard phenol-chloroform extraction method [59]. The experiment was approved by the Institute Animal Ethics Advisory Committee (IAEC) ICAR-Directorate of Poultry Research, Hyderabad, India.

**Fig. 1.**
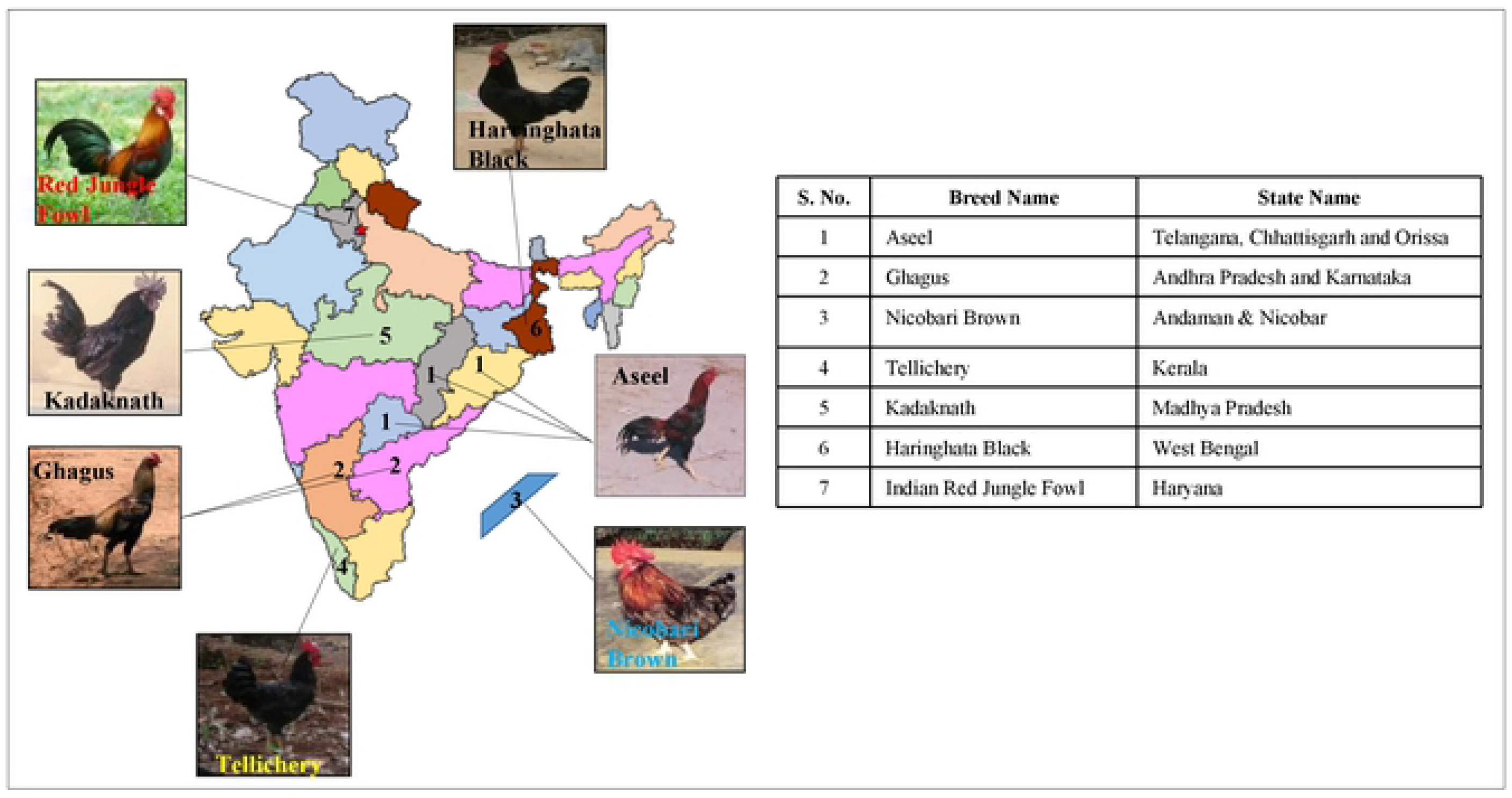
Geographic distribution of Indian native chicken breeds used in the current study.

**Table 1.**
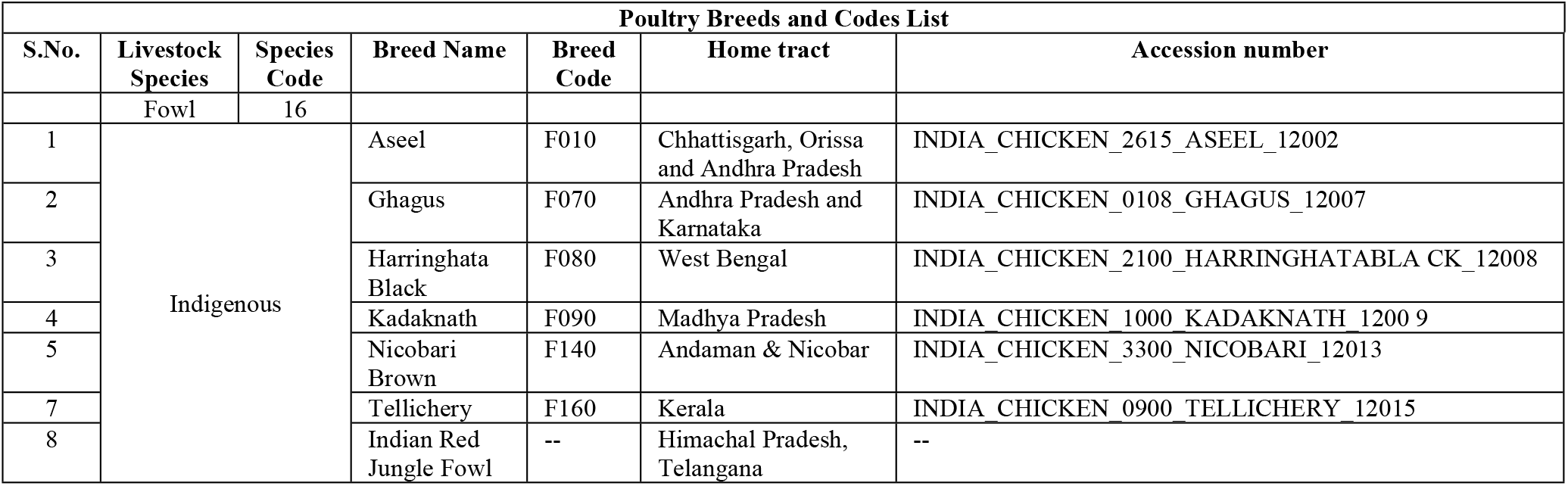
Information for Indian native chicken breeds used in the current study.

**Table 2.** Characteristic features of Indian native chicken breeds used in the current study.

| S.No. | Breed Name | Weight (Avg Kg) |  | Plumage Type | Plumage Pattern | Plumage Colour | Comb Type | Skin Colour | Shank Colour | Egg Shell Colour | Visible Character | Principal use |
| --- | --- | --- | --- | --- | --- | --- | --- | --- | --- | --- | --- | --- |
|  |  | Male | Female |  |  |  |  |  |  |  |  |  |
| 1 | Aseel | 4 | 2.59 | Normal | Patchy | Red, Black | Pea | Yellow | Yellow | Brown | Small but firmly set comb. Bright red ear lobes. Long and slender face devoid of feathers. The general feathering is close, scanty and almost absent on the breast. The plumage has practically no fluff and the feathers are tough. | Socio-Cultural-Game/Fighting; Food-meat |
| 2 | Ghagus | 2.16 | 1.43 | Normal | Patchy | Brown | Pea or single | White | Yellow | Light Brown | Cocks have shining bluish black feathers on breast, tail and thighs. Neck is covered with golden feathers. Throat in some cases is loose and hanging. Wattles are small and red in colour. Ear lobes are mostly red. | Food - meat, eggs |
| 3 | Harringhata Black | 1.284 | 1.121 | Normal | Self-Black | Black | Single | White | Grey | Light Brown | -- | Food-Meat, eggs |
| 4 | Kadaknath | 1.6 | 1.125 | Normal | -- | Ranges from silver to gold spangled to blue black | -- | Dark grey | Grey | Light Brown | The colour of day old chicks is bluish to black with irregular dark stripes over the back. In the adults, comb, wattles and tongue are purple. The shining blue tinge of the ear lobes adds to its unique features. | Food - meat; Socio-cultural - religious ceremonies |
| 5 | Nicobari Brown | 1.8 | 1.3 | Normal | Solid | Black | Single | Dark grey | Grey | Light Brown | The birds are short legged. Shank length at 10 weeks of age varies from 3.50 to 3.85 cm. | Food - Eggs and Meat |
| 6 | Tellichery | 1.62 | 1.24 | Normal | Solid | Black with shining bluish tinge | Single | Grey | Blackish grey | Light Brown | Shining bluish tinge on hackle, back and tail. Comb is red and large in size. It is erect in cocks and drooping on the rear side in hens. Wattles are red in colour. Ear lobe is mostly red in colour. Eye ring is blackish red. Beak is blackish. | Food - meat, eggs |
| 7 | Indian Red Jungle Fowl | 2.04 | 1.36 | Normal | Solid | Bright orange and black | Single | White | Yellow | White | Feathered shank, bunch of feather on head (crown structure) in about 18 % birds | Egg, meat and socio-cultural |

### 2.2. Primer designing

Primer designing, library preparation, and sequencing were performed at Genotypic Technology’s Genomics facility. Eight primer pairs were designed for covering the entire mitochondrial genome. About 10ng of DNA was taken for PCR to amplify explicit products of 2-3kb in size. All the products were checked using a 1% Agarose gel (S1 Fig.). All products were pooled in equal amounts for sonication utilizing Covaris S220.

### 2.3. DNA template library preparation and sequencing by Ion PGM™ sequencer

Library Preparation was done following the Ion Torrent protocol outlined in the Ion plus fragment library kit (ThermoFisher Scientific, USA, # 4471252). About 500ng of fragmented and cleaned DNA was taken for library preparation. End-repair and adapter ligation were done and the samples were barcoded in this progression. The samples were cleaned utilizing AMPure XP beads. Samples were size-chosen utilizing a 2% low melting agarose gel. The gel-purified samples were amplified for the enrichment of adapter-ligated fragments as per the protocol. The amplified products were cleaned using AMPure XP beads and quantified using Qubit Fluorometer and then run on Bioanalyzer High sensitivity DNA Assay to assess the quality of the library. The purified libraries were then used to prepare clonally amplified templated Ion Sphere™ Particles (ISPs) for sequencing on an Ion PGM™ Chip to obtain the essential data coverage. Sequencing was performed on Ion PGM™ sequencer at Genotypic Technology’s Genomics facility, Bengaluru, India.

### 2.4. Quality control for reads and analysis

The samples were sequenced with Ion PGM Sequencer and analyzed with Torrent suite v 3.6. Once the base-calling was done, the raw reads undergo the in-built process of trimming and filtering to get only the high quality reads. Trimming is done to evacuate any undesired base calls, adapter sequences, and lower quality reads at 3’ end of the reads. Filtering at the next step evacuates reads assessed to contain the low-quality base calls. These two steps ensure the reads which taken further are of high quality and the clean reads were used for the subsequent analyses. The raw reads obtained are aligned to the reference Gallus gallus mitochondria (NC_001323.1) with the TMAP algorithm. The variants were detected using the inbuilt plugin Variant caller (V4.0) of TS.

### 2.5. Validation of mtDNA genes with gene specific primers

The mtDNA gene-specific primers were designed by using IDT oligo analyzer software (Table 3). The mtDNA genes were amplified by using the Prima-96™ Thermal Cycler (HIMEDIA). PCR amplification was performed in a 25 μl volume containing 1 μl of DNA template, 2.5 μl of 10× PCR buffer with Mg^2+^, 2.5 μl of dNTP mixture (2.5 mM each), 1.25 μl of each primer (10 μM), 0.5 μl of Taq DNA polymerase (5 U/μl) and 16 μl of nuclease-free water. The PCR conditions were as follows: 95°C for 10 min; followed by 35 cycles of 94°C for 30 s, 53/58°C for 30 s and 72°C for 45 s; and a final extension at 72°C for 10 min. The PCR products were detected on 2% agarose gel electrophoresis.

**Table 3.**
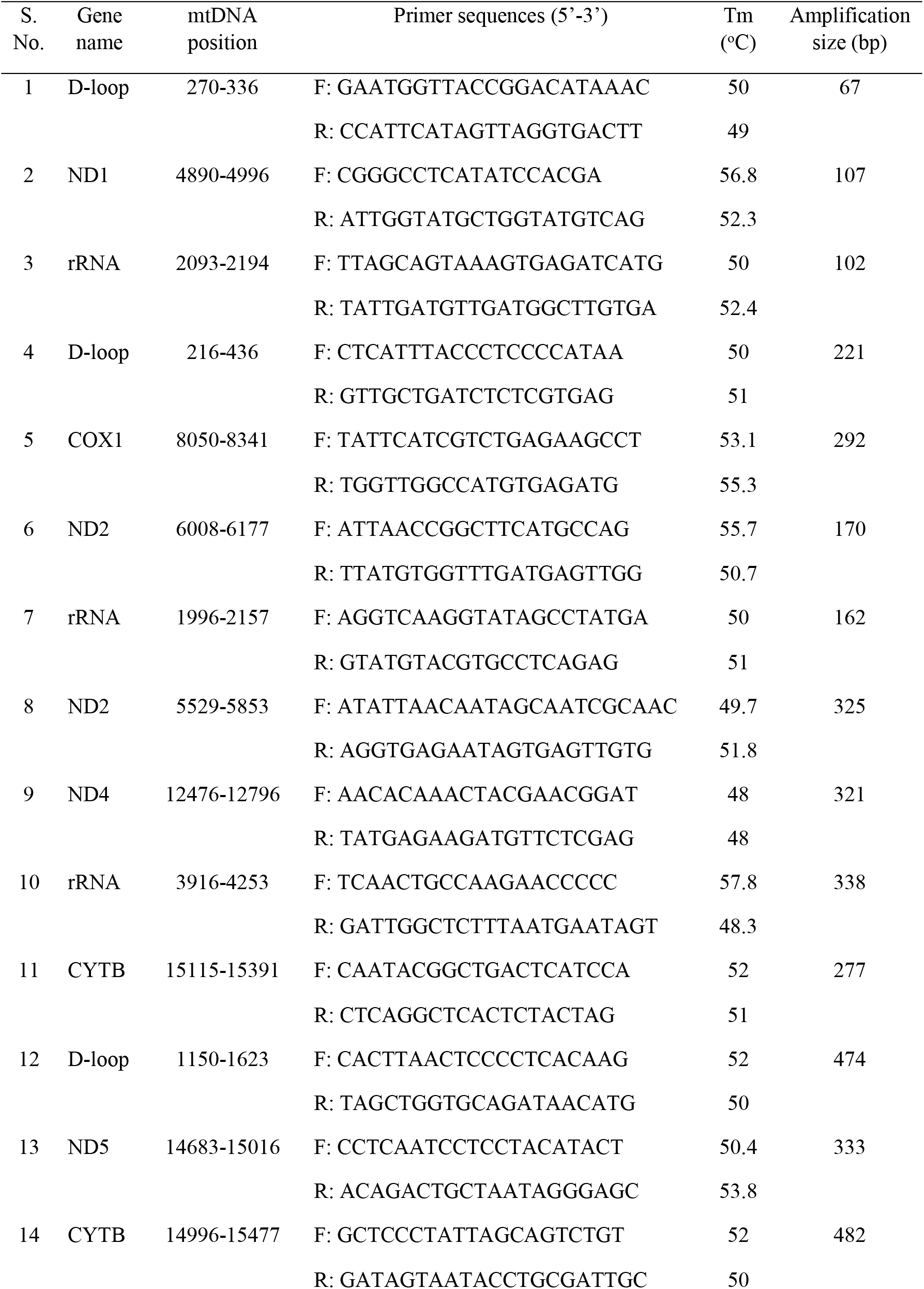
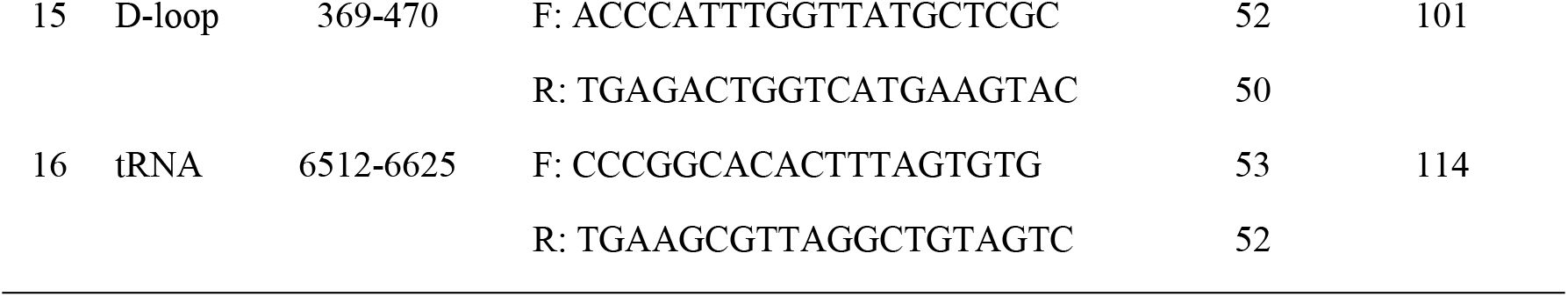
The PCR primers for segmental amplification of chicken mtDNA genes.

### 2.6. Phylogenetic and molecular evolution analysis

To investigate the evolutionary relationships, a phylogenetic examination was performed using the total mitochondrial DNA and D-loop region sequences of seven Indian native chicken breeds along with twenty-two Asian native chicken breeds. Each of the sequence datasets was aligned by Clustal X and analyzed by neighbor-joining (N-J) in MEGA 10.0, and bootstrap analysis was performed with 1000 replications [51–53]. For building a neighbor-joining phylogenetic tree, Kimura’s two parameters model was used for calculating the genetic distance of the haplotypes.

### 2.7. Haplotypes Diversity

Network analysis was used for haplotype diversity illustrated by using NETWORK 10.1 [50]. In chicken, population indices include several segregation sites (*S*), number of haplotypes (*H*), haplotype diversity (Hd), and nucleotide diversity (*π*), determined by the mtDNA D-loop sequences diversity and elucidate the sequence polymorphism and the content of genetic variability [54]. The DnaSP software version 10.1 was used for analysis and the alignment gaps arising from a deletion event were excluded from the calculations [55]. Between two sequences, the average number of nucleotides differences per site known as nucleotide diversity (*π*) is defined as *π* = *n*/ (*n* −1) Σ*xixjπij* or *π* = Σ*πij/nc* [*n*=number of DNA sequences examined; *xi* and *xj*=frequencies of the *i*th and *j*th type of DNA sequences; *πij*=proportion of nucleotides in the respective types of DNA sequences; *nc*=total number of sequence comparisons] [54]. According to Nei formula, the average heterozygosity or haplotype diversity, *h*, is defined [*h* = 2*n* (1 − Σ*xi*2)/ (2*n* − 1); *xi*=frequency of haplotype and *n*=sample size] [54]. The gene or haplotype frequencies were used for assessing the level of genetic differentiation among the population. The *F*st and *N*st significant tests by Arlequin software version 2.000 were used to explore the population’s genetic structure [56–58].

## 3. Results

The proposed whole mtDNA sequence analysis effectively be used to characterize the seven Indian indigenous chickens along with twenty-two Asian native chickens. The blood samples were collected and mtDNA was extracted from seven Indian native chicken breeds, they are Aseel, Ghagus, Nicobari Brown, Nicobari Black, Tellichery, Kadaknath, Haringhata Black, and Red Jungle Fowl. We successfully amplified mtDNA using an amplification method and NGS sequencing was done by using Ion PGM™ sequencer at Genotypic Technology’s Genomics facility. We obtained 757073, 89859, 105503, 50617, 15764, 264309, and 36685 sequencing reads from eight Indian native chickens, respectively (Table 4; Table S1). The assembled complete mitochondrial genomes for seven Indian native chickens have been submitted to Genbank under accession numbers, KP211418.1, KP211419.1, KP211422.1, KP211424.1, KP211425.1, KP211420.1, and KP211423.1 for Aseel, Ghagus, Nicobari Brown, Tellichery, Kadaknath, Haringhata Black, and Indian Red Jungle Fowl, respectively.

**Table 4.** Sequencing reads obtained from seven Indian native breeds.

|  | Aseel | Ghagus | Nicobari brown | Tellicherry | Kadaknath | Haringhata black | Indian Red Jungle Fowl |
| --- | --- | --- | --- | --- | --- | --- | --- |
| Total number of reads | 757073 | 89859 | 105503 | 50617 | 15764 | 264309 | 36685 |
| Number of mapped reads | 734474 | 87336 | 103692 | 49407 | 15224 | 257435 | 35693 |
| Average base coverage depth | 6303 | 593 | 799 | 387.1 | 90.22 | 1672 | 278.3 |
| Uniformity of base coverage (%) | 84.15 | 76.81 | 81.78 | 61.88 | 68.34 | 88.98 | 71.18 |
| Genome base coverage at 1x (%) | 100.00 | 100 | 100 | 100 | 99.68 | 100 | 100 |
| Genome base coverage at 20x (%) | 100.00 | 99.87 | 99.57 | 91.16 | 65.31 | 100 | 91.12 |

The complete length of mtDNA of Indian local chickens was 16,775bp. Like most different vertebrates, it contains the common structure, including 2 rRNA genes, 22 tRNA genes, 12 protein-coding genes, and 1 D-loop region [60,61]. The four nucleotides (i.e. A, T, G, C) overall composition of assessed mtDNA was 30.26%, 23.76%, 32.48%, and 13.50%, in the order G > A > T > C, respectively. The inception codons for all the coding proteins are ATG, aside from COX1 which is GTG (Table 5). The heavy (H) strand of mtDNA encoded all the mtDNA genes and light (L) strand encoded four sorts of tRNA genes and ND6 genes. Every one of these genes have15 spaces in the length of 2-227bp and have 3 overlaps in the length of 1-8bp. These genes had three sorts of termination codons, including TAA, TAG, and “TGA”. The “T– –” is the 5’ terminal of the adjoining gene [62]. The lengths of the two rRNA genes were 976bp and 1621bp. Among 12 protein-coding genes, the longest one was the ND5 gene (1818 bp) and the most limited one was the ND4L gene (297bp). Like different types of chicken, 4 tRNA genes were circulated in protein-coding genes, vary from 66 to 135bp in size [63–66]. The D-loop region was situated among ND6 and rRNA with a length of 1227bp (Table 5; Table S2).

**Table 5.** Organization of the mitochondrial genome of eight native Indian chickens.

| Gene name | Position |  | Size | Codon |  | Anticodon | Strand | Space/Overlap+ |
| --- | --- | --- | --- | --- | --- | --- | --- | --- |
|  | Start | End |  | Start | Stop |  |  |  |
| D-loop | 1 | 1227 | 1227 |  |  |  |  |  |
| rRNA | 1297 | 2272 | 976 |  |  |  | H | 70 |
| rRNA | 2346 | 3966 | 1621 |  |  |  | H | 74 |
| ND1 | 4050 | 5024 | 975 | ATG | TAA |  | H | 84 |
| ND2 | 5241 | 6281 | 1041 | ATG | TAG |  | H | 217 |
| tRNA-Cys | 6508 | 6573 | 66 |  |  | GCA | L | 227 |
| COX1 | 6645 | 8132 | 1488 | ATG | T-- |  | H | 72 |
| tRNA-Ser | 8124 | 8258 | 135 |  |  | UGA/GCU | L | -8 |
| tRNA-Asp | 8261 | 8329 | 69 |  |  | GUC | L | 3 |
| COX2 | 8331 | 9014 | 684 | ATG | TAA |  | H | 2 |
| ATP6 | 9240 | 9923 | 884 |  |  |  | H | 226 |
| COX3 | 9923 | 10706 | 784 | ATG | T-- |  | H | -1 |
| ND3 | 10776 | 11126 | 351 | ATG | TAA |  | H | 70 |
| ND4L | 11196 | 11492 | 297 | ATG | TAA |  | H | 70 |
| ND4 | 11486 | 12863 | 1378 | ATG | T-- |  | H | -6 |
| ND5 | 13071 | 14888 | 1818 | ATG | TAA |  | H | 208 |
| **CYTB** | 14893 | 16035 | 1143 | ATG | TAA |  | H | 5 |
| **tRNA-Pro** | 16108 | 16177 | 70 |  |  | UGG | L | 73 |
| **ND6** | 16184 | 16705 | 522 | ATG | TAA |  | L | 7 |

T– means incomplete termination codon. +Negative numbers indicate overlapping nucleotides.

### 3.1. Pattern of mtDNA D-Loop Variability

The 1227bp mtDNA D-loop fluctuation design uncovered high variations between nucleotides 235 and 1209. In seven Indian local chicken breeds, 8 haplotypes were related to variation at 28 destinations, and 8.33% of them were polymorphic (Fig. 2). The total arrangement uncovered exceptionally high changeability in the mtDNA D-loop region between 164-360 bases; this variation comprises 23.5% of the 7 successions. This rate is amazingly high contrasted with the other local chicken varieties 5.54-7.05% [24,47]. This high pace of mtDNA D-loop variation might be credited to the relocation of birds all through the nation and diverse topographical districts in India. The base composition of the Indian native chicken breeds mtDNA D-loop shows that A+T grouping content establishes 60.39% while G+C was 39.61% [67].

**Fig. 2.**
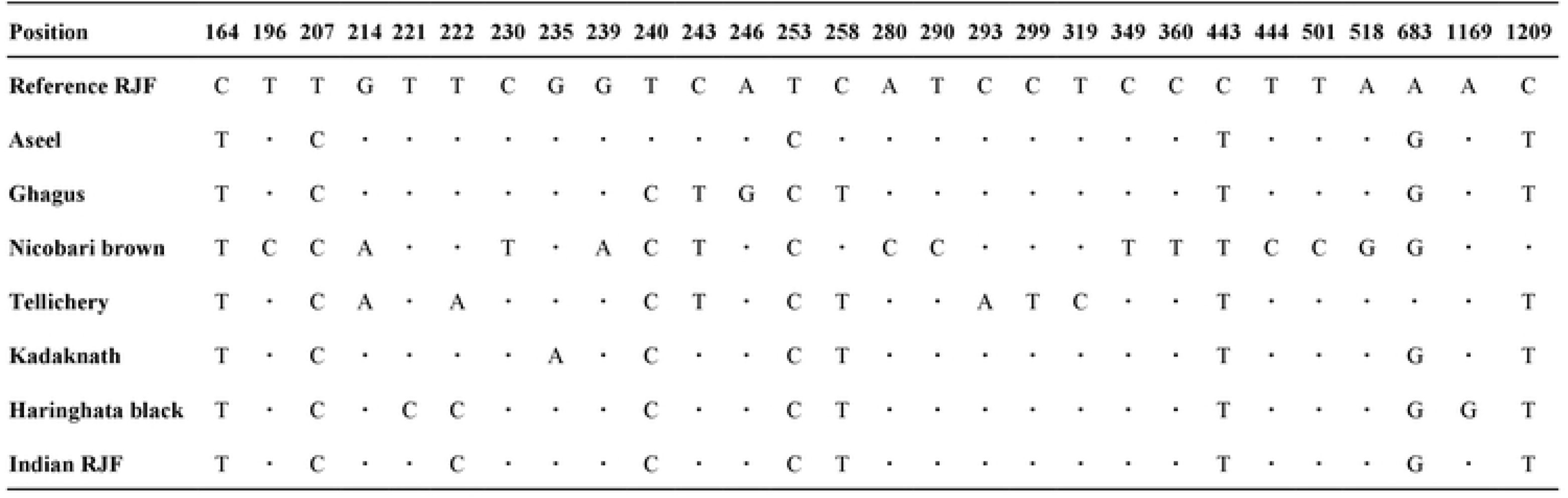
Pattern of mtDNA D-loop variability. Nucleotide polymorphisms observed in D-loop region of seven Indian native chicken sequences. Vertically oriented numbers indicate the site position and the sequences shown are only the variable sites. Dots (.) indicate identity with the reference sequence and different base letters denote substitution.

### 3.2. Sequence variation and haplotype distribution

Multiple sequence alignment was performed for seven Indian native chicken varieties and recognized eight haplotypes. The alignment of D-loop sequences was finished with a gallus reference sequence (NC_001323.1) utilizing Clustal-X. The accompanying domains and motifs were watched, at the 5’ end of the D-loop. Two units of invariant tetradecamer 5’-AACTATGAATGGTT-3’ were distinguished at positions 264 to 277 and 325 to 338. An interfered with thymine string (TTTTATTTTTTAA) was observed as conserved in all the individuals contemplated. There was likewise an intrusion of poly-C sequence (5’-CCCCCCCTTTCCCC-3’) which is generally conserved and downstream to this there is a preserved sequence known as poly-G (5’-AGGGGGGGT-3’). Two moderated 5’-TACAT-3’ and 5’-TATAT-3’ were likewise found in all individuals. There are nine TATAT motifs and four TACAT found inside the D-loop and were thus, conserved. The initial 163 base sets adjoining tRNA Glu were seen as exceptionally conserved in all individuals. The nucleotide replacements found in the 9 variable haplotypes contained one G/T and two C/A transversions and the rest were all changes of which four were A/G substitutions and seven were C/T substitutions. This exhibits a solid predisposition towards transition. The C/T substitutions are more normal than A/G substitution (Table 6).

**Table 6.**
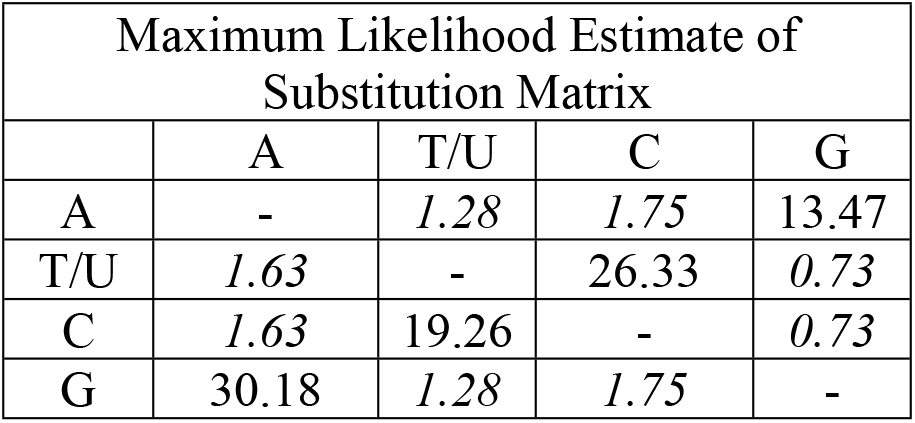
Each entry is the probability of substitution (r) from one base (row) to another base (column). Substitution pattern and rates were estimated under the Tamura-Nei (1993) model [96]. Rates of different transitional substitutions are shown in bold and those of transversionsal substitutions are shown in italics. Relative values of instantaneous r should be considered when evaluating them. For simplicity, sum of r values is made equal to 100, The nucleotide frequencies are A = 30.26%, T/U = 23.76%, C = 32.48%, and G = 13.50%. For estimating ML values, a tree topology was automatically computed. The maximum Log likelihood for this computation was −23504.716. This analysis involved 9 nucleotide sequences. Codon positions included were 1st+2nd+3rd+Noncoding. There were a total of 16775 positions in the final dataset. Evolutionary analyses were conducted in MEGA X [97].

### 3.3. Genetic distance among seven Indian native chicken breeds

The genetic distances within and between the seven Indian native chicken breeds were analyzed by using the DnaSP program 10.1 (Table 7). The genetic distance within the seven chicken breeds was 0.000655~16.173895. Indian Red Jungle Fowl chickens had the most noteworthy within-breed genetic distance, while Ghagus chickens had the least. Between the breeds, the genetic distance estimates went from 0.000655 to 16.188613. The genetic distance between the breeds was most noteworthy for Indian Red Jungle Fowl and Tellicherry chickens, while it was least for Ghagus and Aseel chickens. The incredible number of variants and SNPs were recognized in Nicobari Brown and a minimal number of variants and SNPs are distinguished in Tellicherry and Kadaknath, respectively. The most number of INDELs were distinguished in Kadaknath and the least number of INDELs were recognized in Tellicherry (Table 8).

**Table 7.**
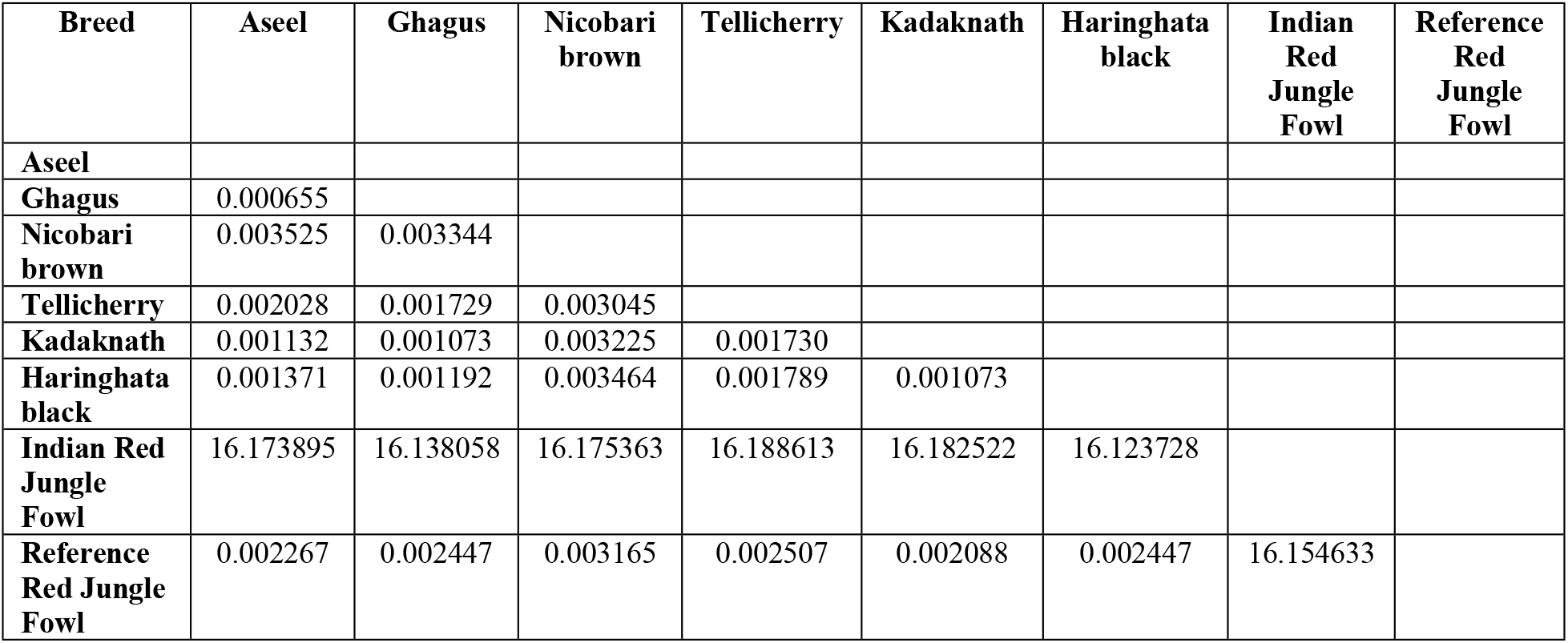
Genetic distance between each pair among the seven Indian native chicken breeds.

**Table 8.**
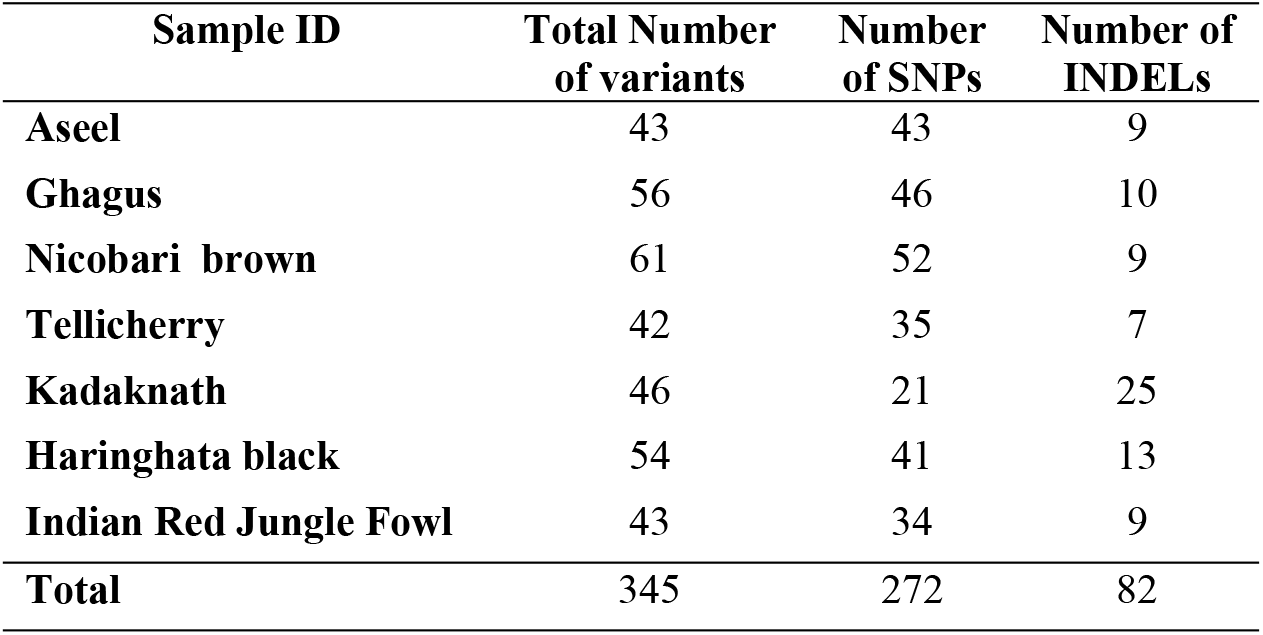
The number of variants, SNPs and INDELs identified in seven Indian chicken mitochondrial genome.

| Sample ID | Total Number<br>of variants | Number<br>of SNPs | Number of<br>INDELs |
| --- | --- | --- | --- |
| **Aseel** | 43 | 43 | 9 |
| **Ghagus** | 56 | 46 | 10 |
| **Nicobari brown** | 61 | 52 | 9 |
| **Tellicherry** | 42 | 35 | 7 |
| **Kadaknath** | 46 | 21 | 25 |
| **Haringhata black** | 54 | 41 | 13 |
| **Indian Red Jungle Fowl** | 43 | 34 | 9 |
| **Total** | 345 | 272 | 82 |

### 3.4. Phylogenetic Analysis of the Haplotypes

An N-J tree indicated that the examined Asian native breeds and Indian local varieties are separated into six significant clades: A, B, C, D, E, and F, besides, the Indian native breeds are situated at the base of the tree (Fig. 3). This N-J tree produced from the total mitochondrial DNA of twenty-two Asian native breeds and seven Indian local varieties has comparative geographies. The clades A to E shared the Asian native breeds but clade F shared only Indian native breeds. Along these lines, our outcomes affirmed that Kadaknath-Haringhata black and Aseel-Ghagus breeds have a nearby hereditary relationship. In addition, the N-J tree generated from the NADH dehydrogenase subunit genes, cytochrome c oxidase subunit genes, mitochondrial encoded ATP synthase membrane subunit 6 gene, cytochrome b gene, and ribosomal RNA genes. The results showed a close hereditary relationship with Aseel-Ghagus and Nicobari brown-Reference RJF (Fig. 5; Fig. 6; Fig. 7; Fig. 8).

**Fig. 3.**
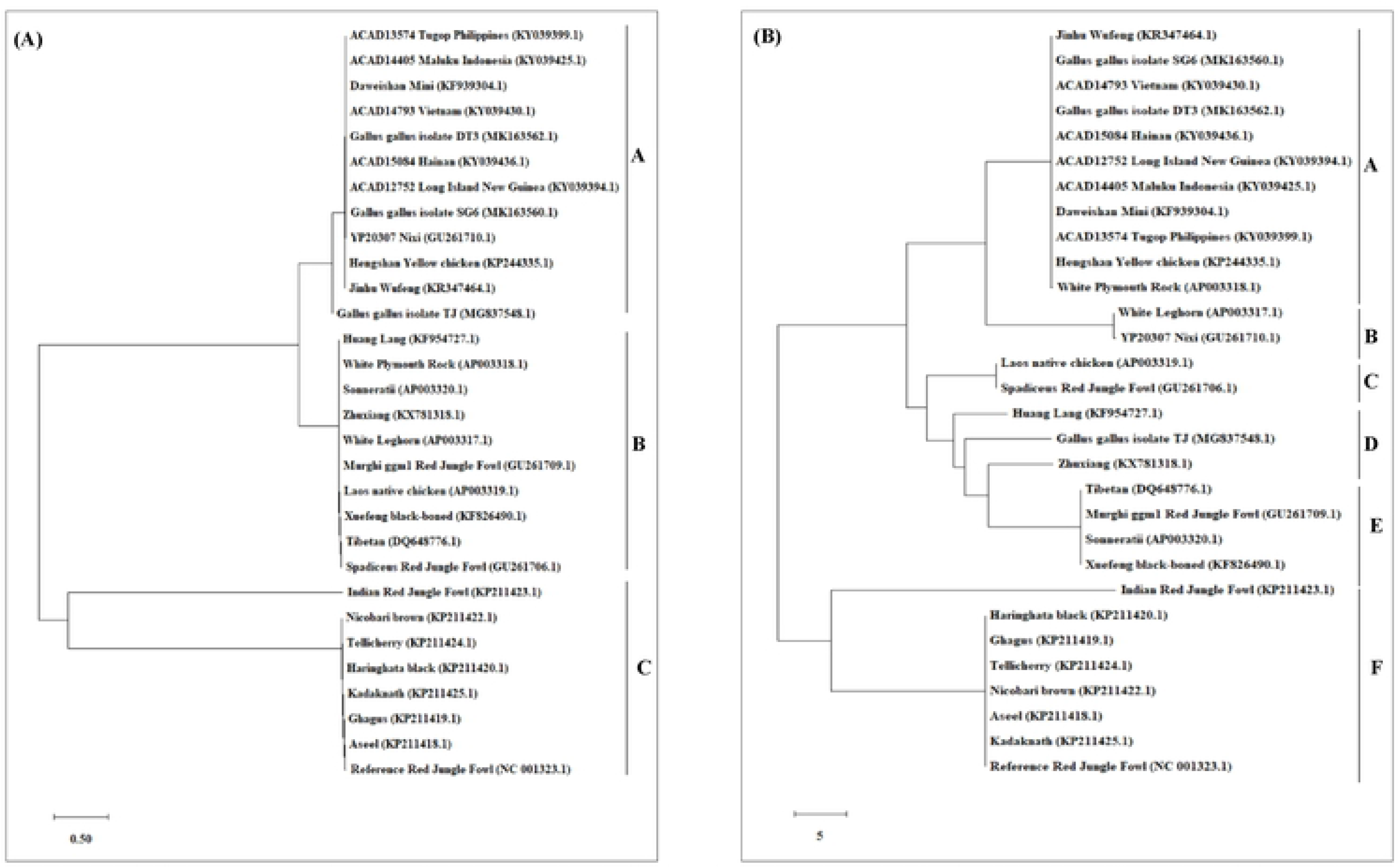
An N-J tree was constructed using MEGA 10.1 software. Phylogenetic analysis based on mtDNA D-loop region (A) and complete mtDNA genome sequences (B) of seven Indian native breads along with reference RJF mtDNA and twenty-two Asian native breeds sequence. The mtDNA genome sequences were obtained from the NGS sequencing and submitted in NCBI (Accession numbers: Aseel-KP211418.1; Ghagus-KP211419.1; Haringhata Black-KP211420.1; Kadaknath-KP211425.1; Nicobari Brown-KP211422.1; Indian Red Jungle Fowl-KP211423.1; Tellichery-KP211424.1). The numbers at the nodes represent the percentage bootstrap values for interior branches after 1000 replications.

### 3.5. Network Analysis

Median-joining networks were drawn for the 8 haplotypes identified from the seven Indian native chickens along with reference Red Jungle Fowl mtDNA, based on the variable characters of the complete alignment using the computer program NETWORK 10.1 [50]. The results showed that the mtDNA sequence of Indian Red Jungle Fowl has the highest frequencies and this haplotype is connected to the frequencies of other haplotypes forming star-like connections. It was also observed that there are mutational links to eight haplotypes with five median vector (mv)∗ separating clades (Fig. 4). The median-joining network examination was completed with the haplotypes from the Indian native breeds. The outcomes show that out of the seven recognized native breeds haplotypes, just two breeds haplotypes i.e. Kadaknath and Haringhata black showed uniqueness and it fell into a different clade and separated from other breeds. Also, median-joining networks were drawn for mtDNA structural genes, such as NADH dehydrogenase subunit genes, cytochrome c oxidase subunit genes, mitochondrial encoded ATP synthase membrane subunit 6 gene, cytochrome b gene, and ribosomal RNA genes. The results showed that three breeds haplotypes i.e. Indian RJF, Kadaknath, and Haringhata black have uniqueness (Fig. 5; Fig. 6; Fig. 7; Fig. 8).

**Fig. 4.**
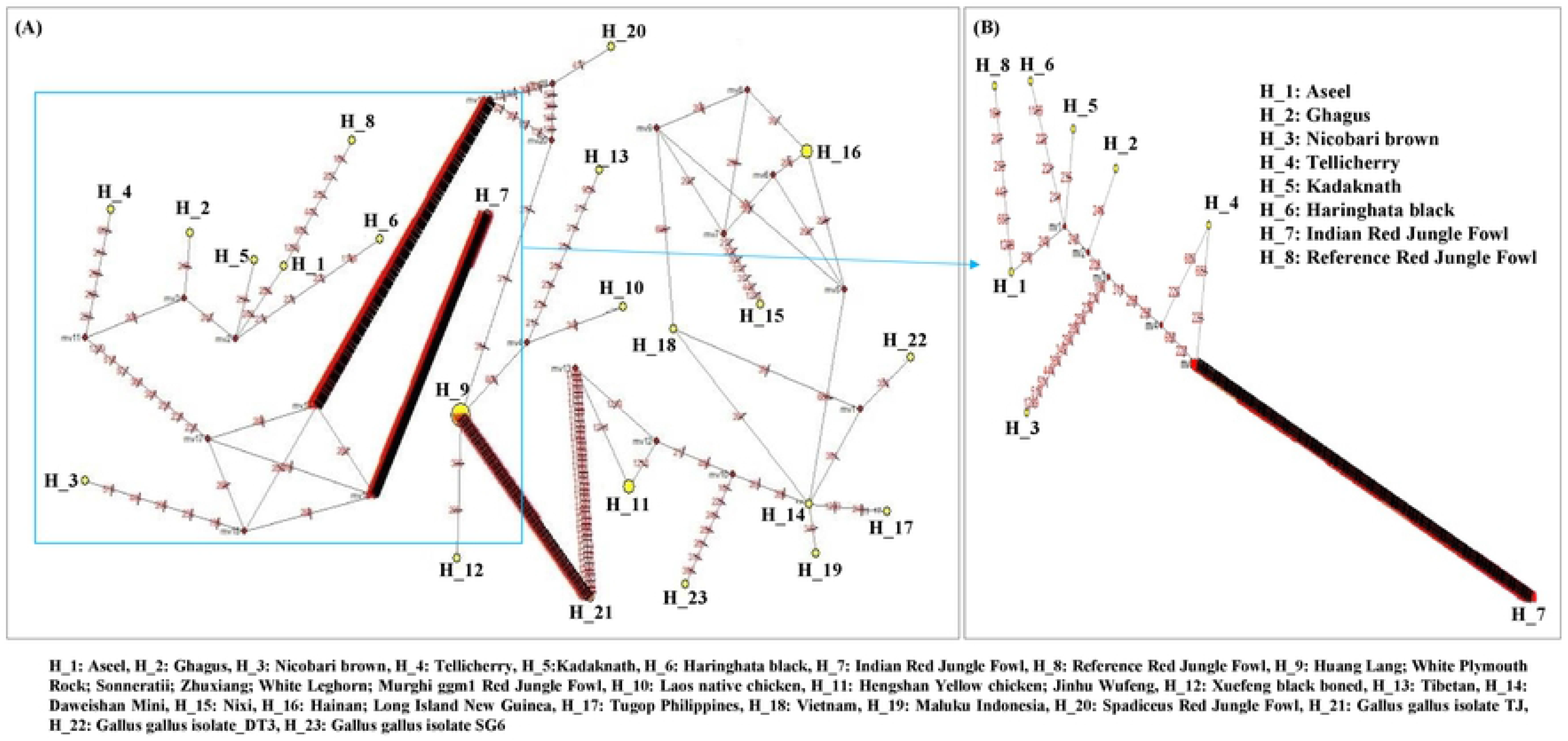
Median-joining network was produced by the network software 10.1 for 8 haplotypes of Indian native chicken breeds along with reference mtDNA based on polymorphic site of the mtDNA D-loop region (A) and complete mtDNA sequences (B). Area of each yellow colour circle is proportional to the frequency of the corresponding haplotype. The pink dots illustrate median vectors (mv) and the numbers on each link line represents mutated positions.

**Fig. 5.**
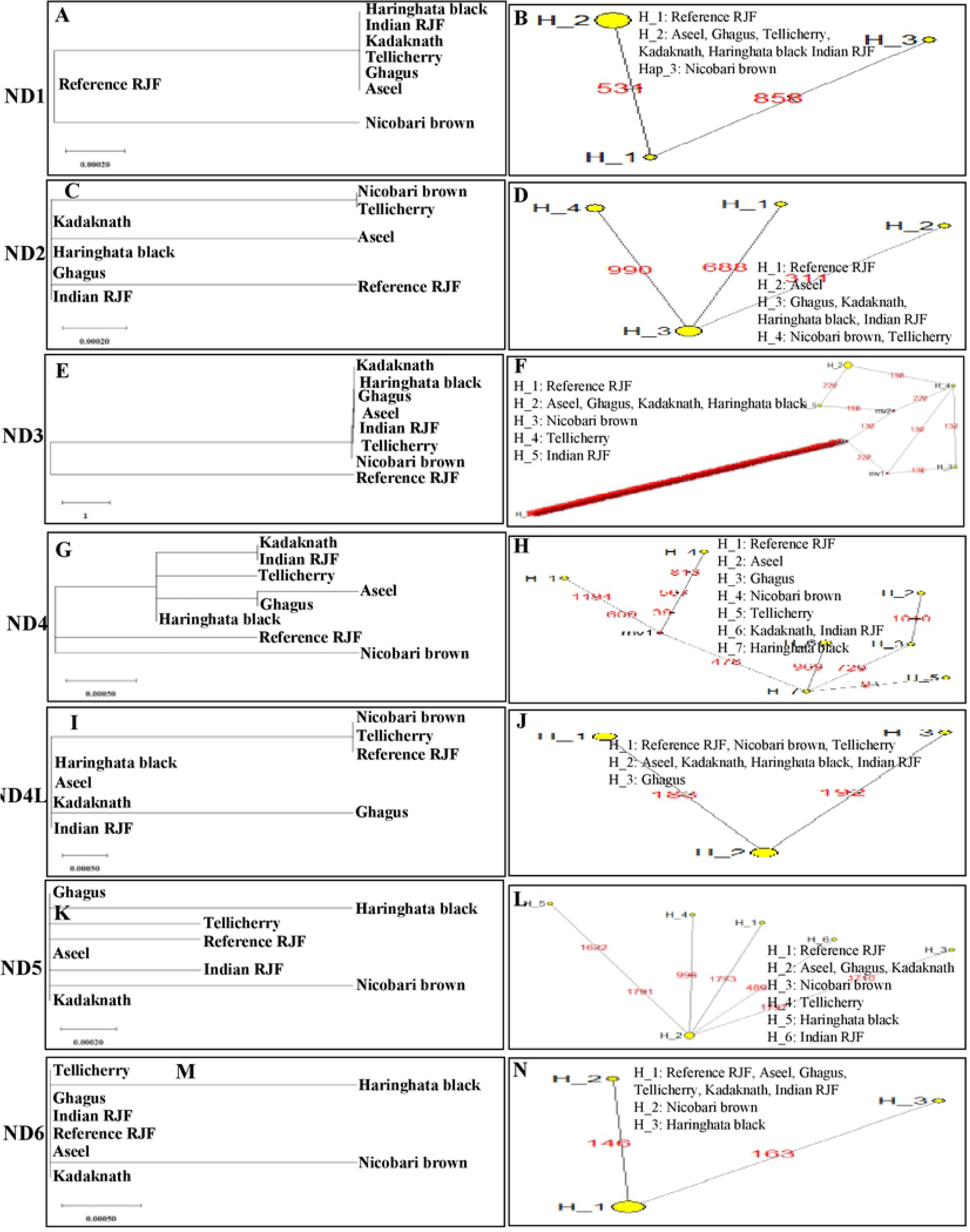
An N-J tree was constructed using MEGA 10.1 software. Phylogenetic analysis (A, C, E, G, I, K, M) based on mtDNA NADH dehydrogenase subunit gene sequences of seven Indian native breeds along with reference RJF. The numbers at the nodes represent the percentage of bootstrap values for interior branches after 1000 replications. Medianjoining network was produced by the network software 10.1 for Indian native chicken breeds along with reference mtDNA based on polymorphic site of the mtDNA NADH dehydrogenase genes (B, D, F, H, J, L, N). The area of each yellow colour circle is proportional to the frequency of the corresponding haplotype. The pink dots illustrate median vectors (mv) and the numbers on each link line represent mutated positions.

**Fig. 6.**
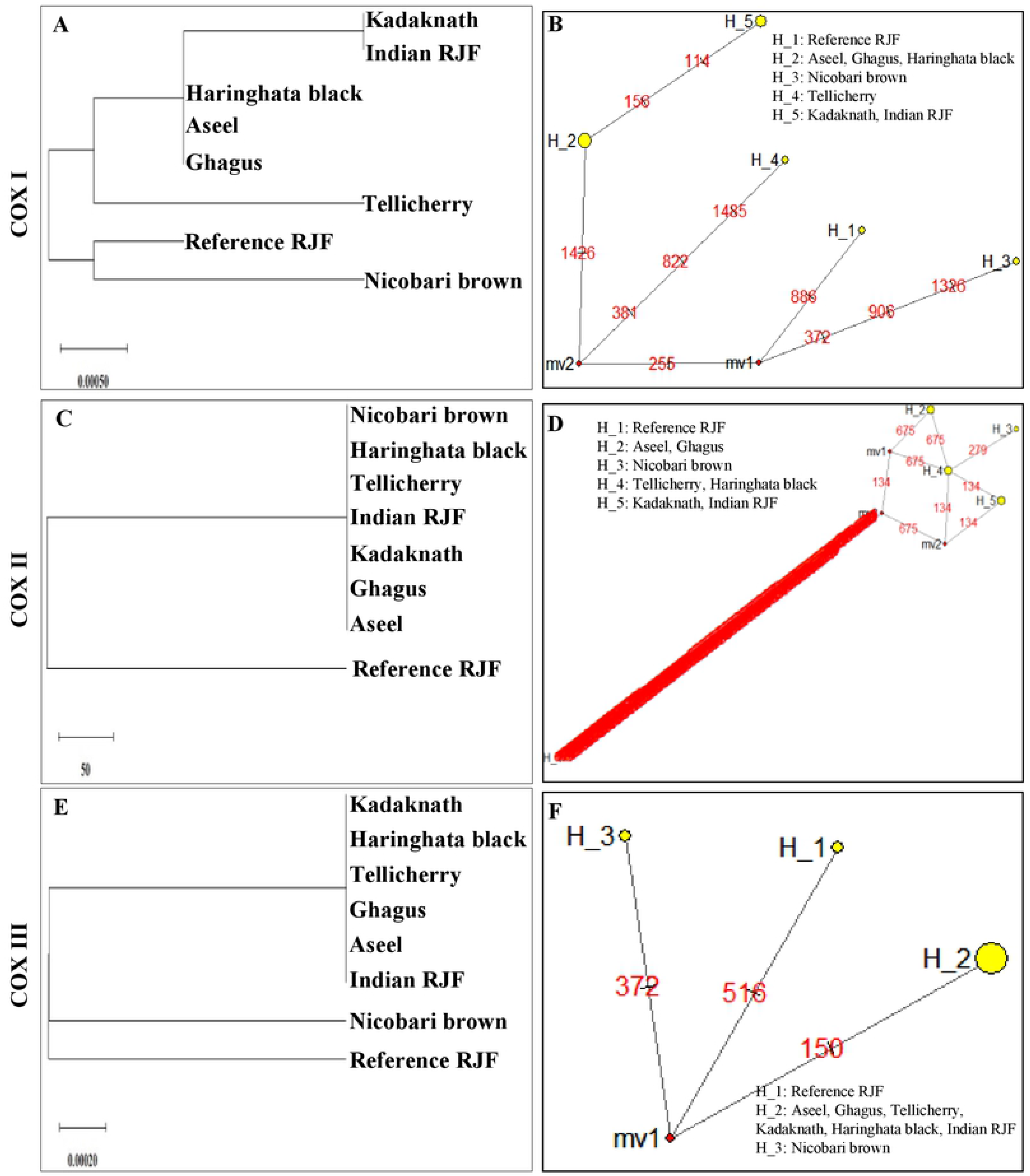
An N-J tree was constructed using MEGA 10.1 software. Phylogenetic analysis (A, C, E) based on mtDNA Cytochrome c oxidase subunit gene sequences of seven Indian native breeds along with reference RJF. The numbers at the nodes represent the percentage of bootstrap values for interior branches after 1000 replications. Median-joining network was produced by the network software 10.1 for Indian native chicken breeds along with reference mtDNA based on polymorphic site of the mtDNA Cytochrome c oxidase subunit genes (B, D, F). The area of each yellow colour circle is proportional to the frequency of the corresponding haplotype. The pink dots illustrate median vectors (mv) and the numbers on each link line represent mutated positions.

**Fig. 7.**
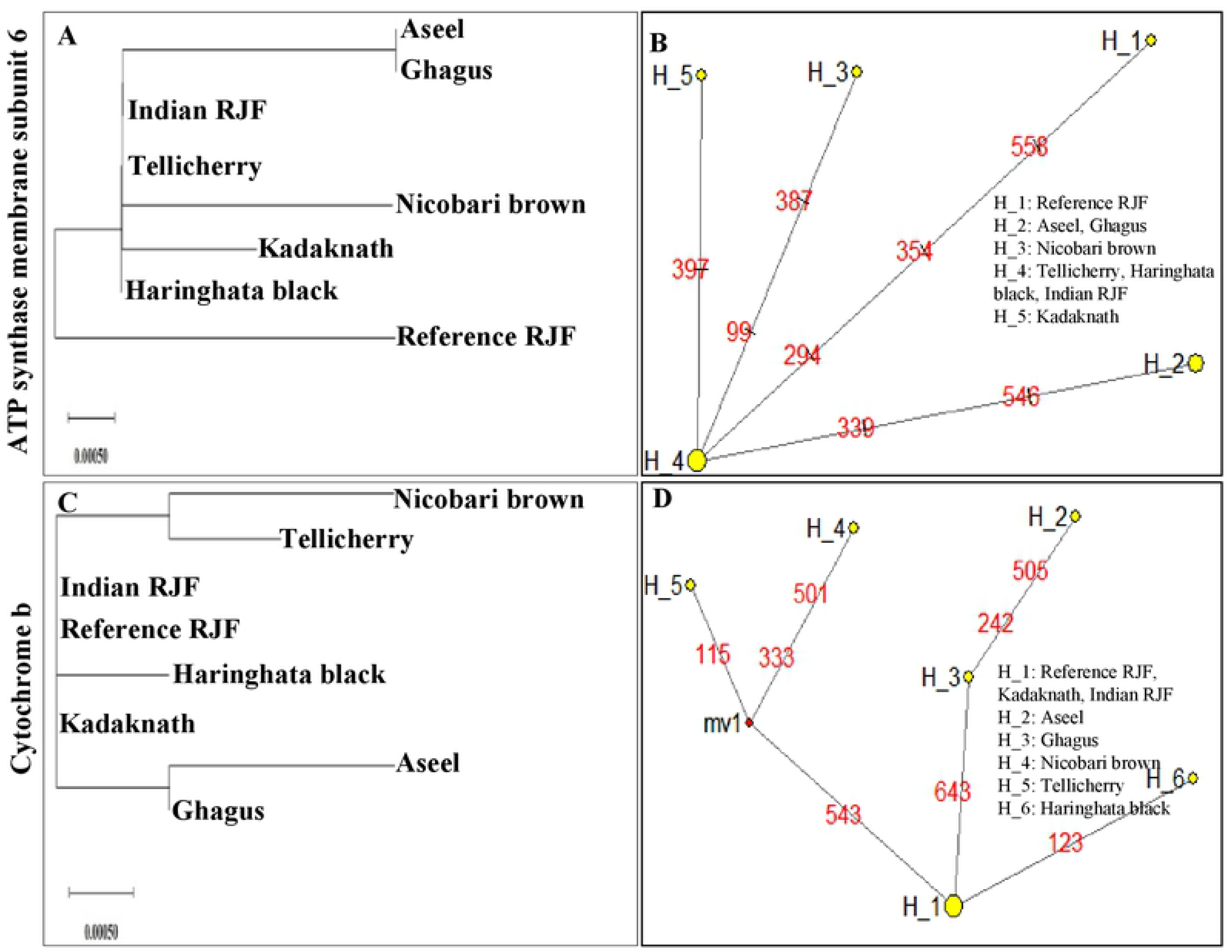
An N-J tree was constructed using MEGA 10.1 software. Phylogenetic analysis (A, C) based on mtDNA ATP synthase membrane subunit 6 and Cytochrome b gene sequences of seven Indian native breeds along with reference RJF. The numbers at the nodes represent the percentage of bootstrap values for interior branches after 1000 replications. Median-joining network was produced by the network software 10.1 for Indian native chicken breeds along with reference mtDNA based on polymorphic site of the mtDNA ATP synthase membrane subunit 6 and Cytochrome b genes (B, D). The area of each yellow colour circle is proportional to the frequency of the corresponding haplotype. The pink dots illustrate median vectors (mv) and the numbers on each link line represent mutated positions.

**Fig. 8.**
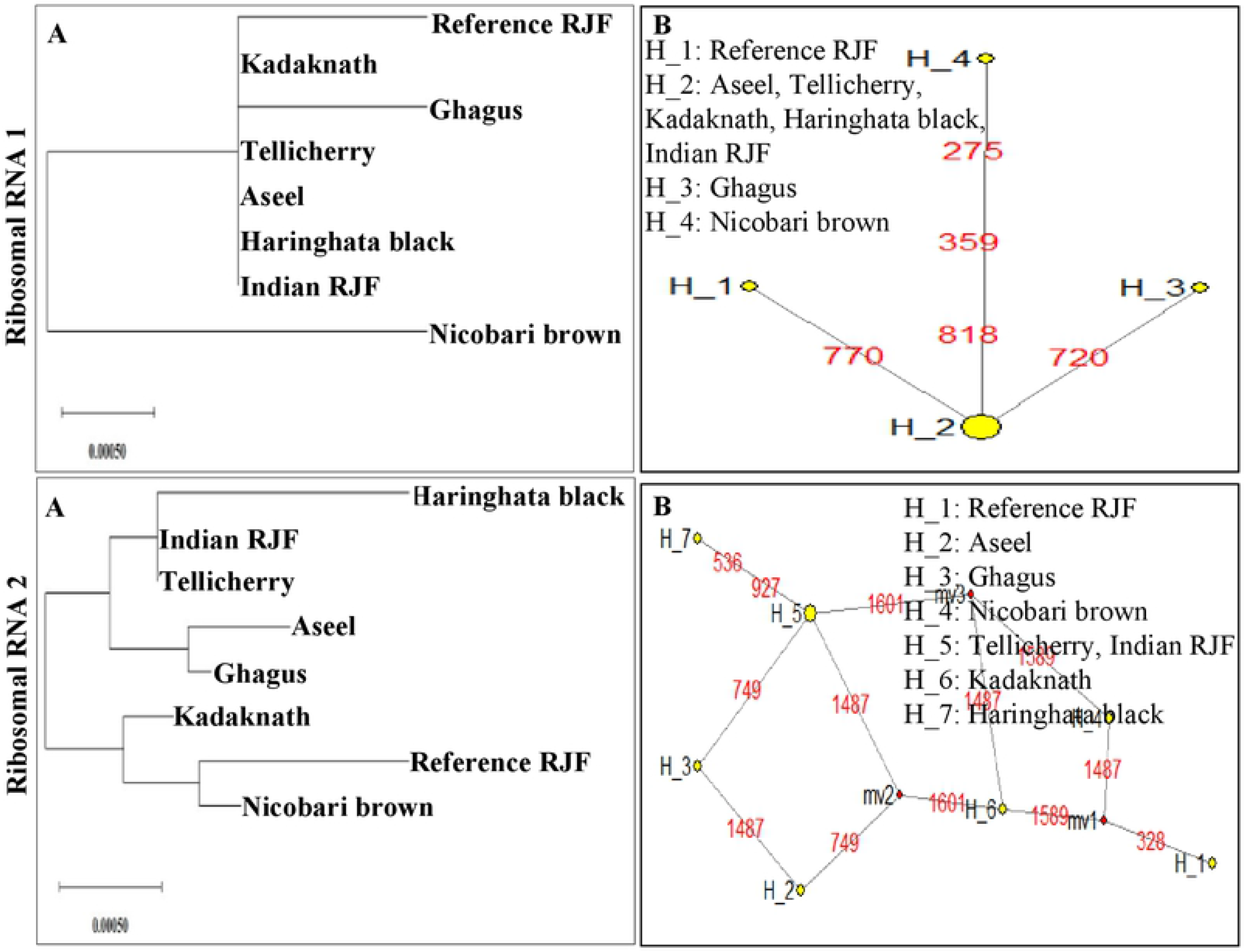
An N-J tree was constructed using MEGA 10.1 software. Phylogenetic analysis (A, C) based on mtDNA Ribosomal RNA 1 and 2 gene sequences of seven Indian native breeds along with reference RJF. The numbers at the nodes represent the percentage of bootstrap values for interior branches after 1000 replications. Median-joining network was produced by the network software 10.1 for Indian native chicken breeds along with reference mtDNA based on polymorphic site of the mtDNA Ribosomal RNA 1 and 2 genes (B, D). The area of each yellow colour circle is proportional to the frequency of the corresponding haplotype. The pink dots illustrate median vectors (mv) and the numbers on each link line represent mutated positions.

### 3.6. The mtDNA genes validation

Genomic DNA was extracted from blood samples of seven Indian native chicken breeds and mtDNA genes were validated by PCR using gene-specific primers. PCR was able to amplify the segmental mtDNA region from all seven native breeds. Each expected PCR amplified product demonstrated the sign band of approximately 67-482 bp in length on 2% agarose gel (Table 3; Fig. 9; Fig. 10; Fig. 11; Fig. 12; Fig. 13). The Aseel specific ND4 primers (12476-12796 region) amplified all the breeds except Ghagus, whereas Aseel rRNA specific primers (3916-4253 region) amplified all the breeds except Ghagus and Haringhata black. The Ghagus specific ND4 primers (11366-11530 region) amplified Indian RJF, Nicobari, Kadaknath, and Ghagus but not amplified Aseel, Haringhata black, and Tellicherry, whereas Ghagus rRNA specific primers (1996-2157 region) amplified all the breeds except Tellicherry. The Kadaknath COX1 specific primers (8050-8341 region) amplified all the breeds except Tellicherry. Such variabilities in amplification of specific fragments occurred due to mutation in the 3’end of the primers as we have designed allele specific primers for amplification of those genes in the specific chicken breeds.

**Fig. 9.**
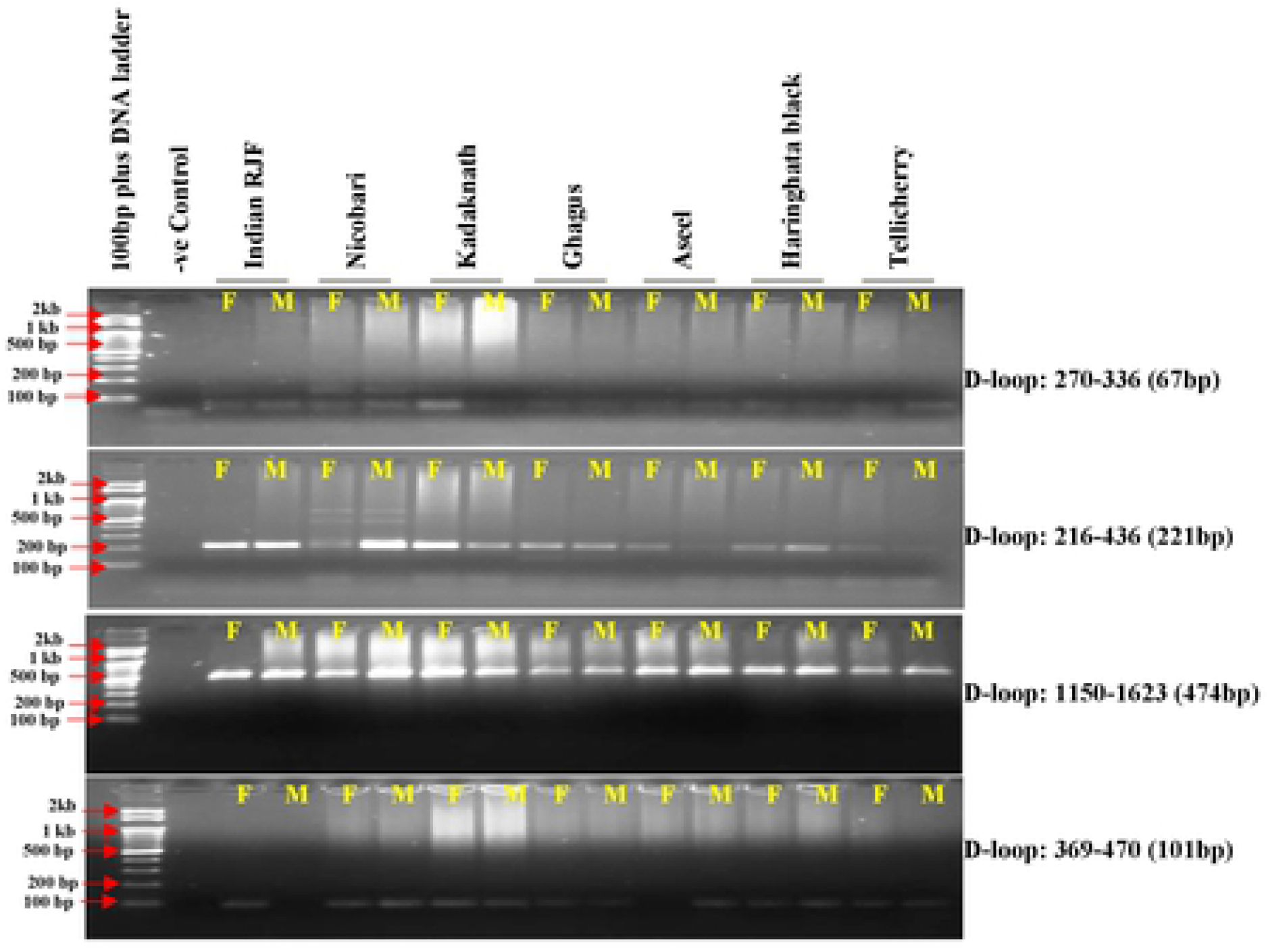
Validation of mtDNA D-loop region with D-loop specific primers.

**Fig. 10.**
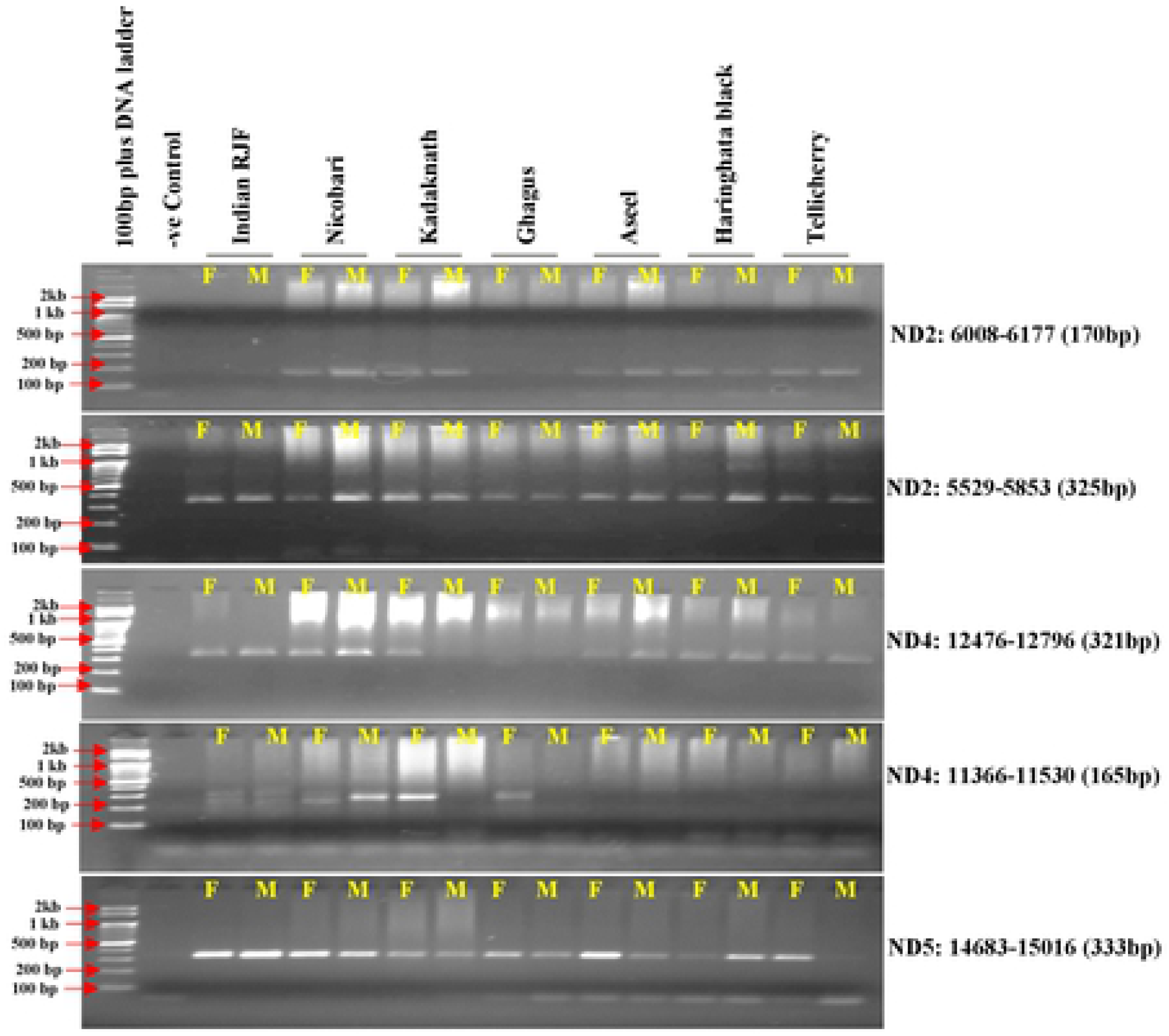
Validation of mtDNA NADH dehydrogenase genes region with gene specific primers.

**Fig. 11.**
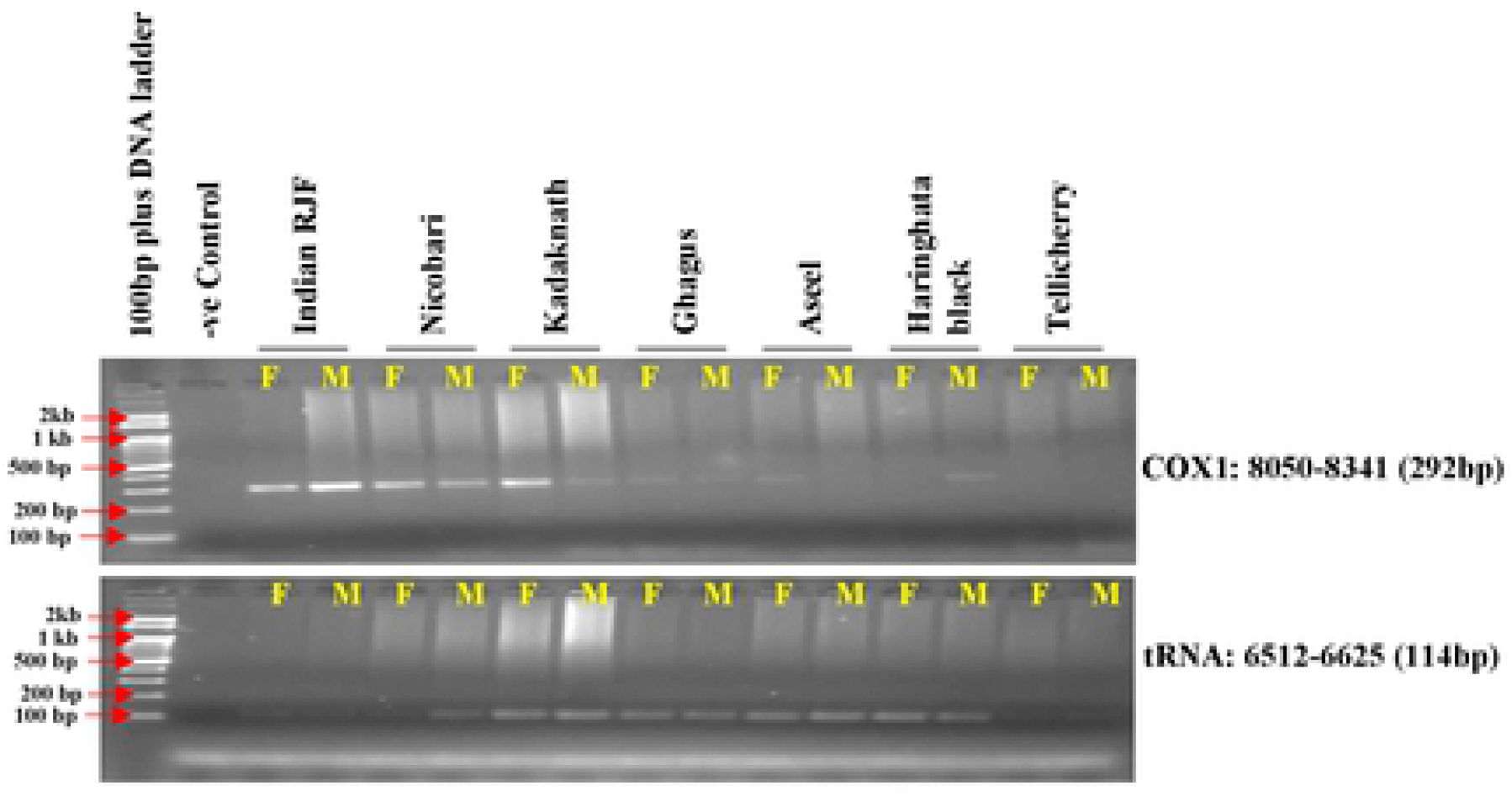
Validation of mtDNA COX1 and tRNA genes region with gene specific primers.

**Fig. 12.**
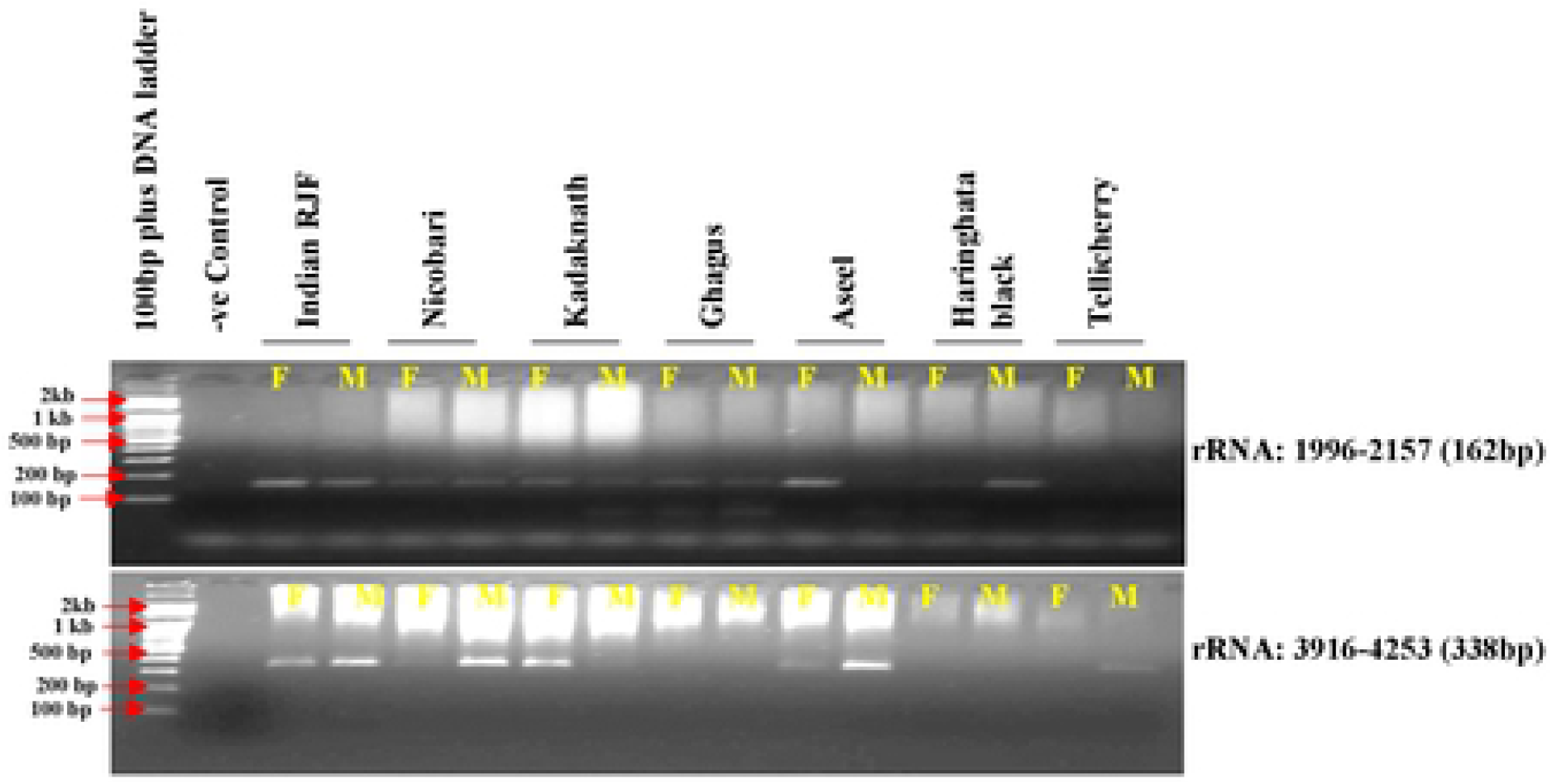
Validation of mtDNA rRNA genes region with gene specific primers.

**Fig. 13.**
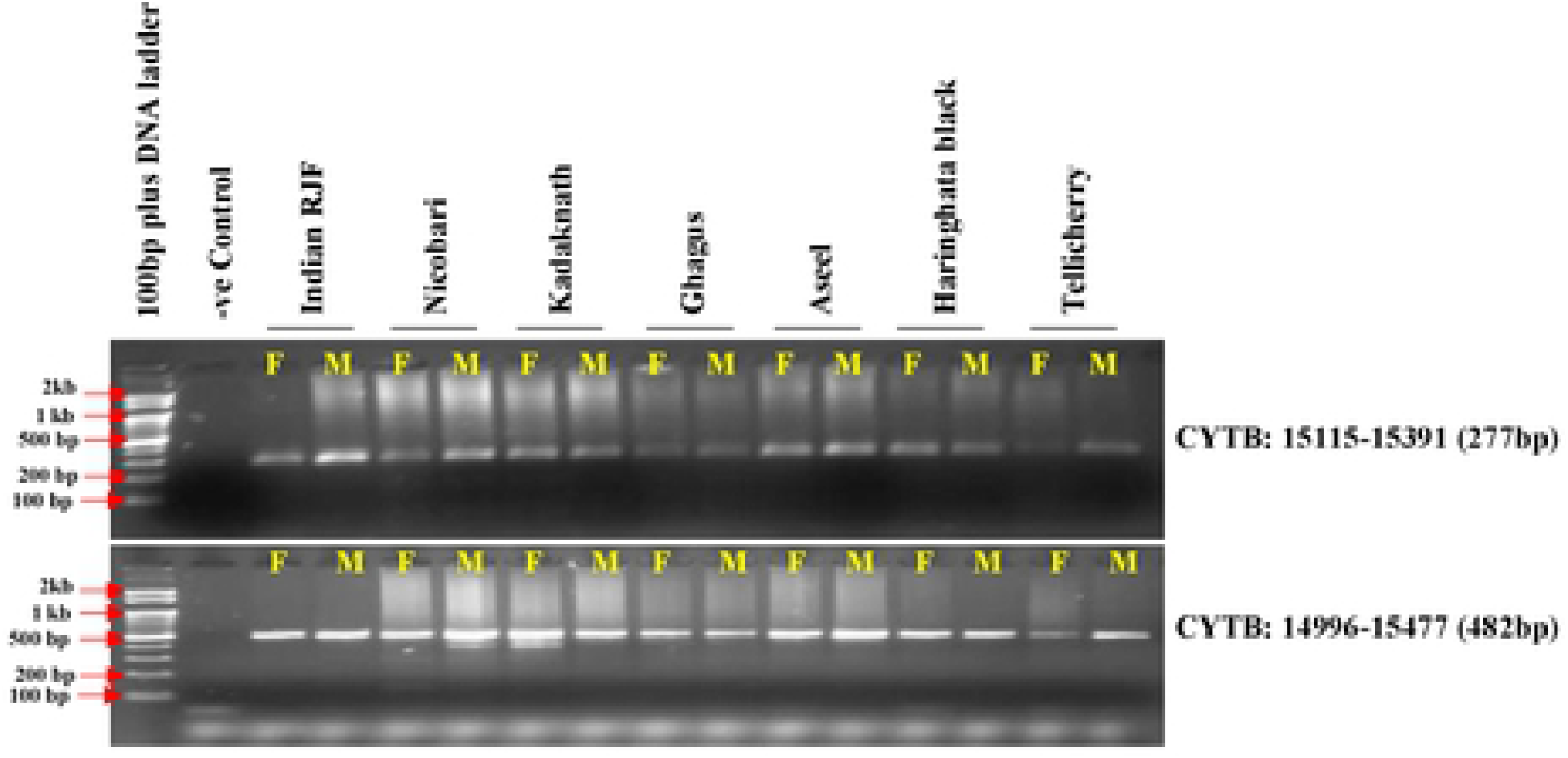
Validation of mtDNA rRNA genes region with gene specific primers.

## 4. Discussion

It is hypothesizing that almost 10,000 years ago the chickens (Gallus gallus domesticus) got domesticated from wild jungle fowls in Southeast Asia [68]. In most of the developing and underdeveloped countries, indigenous/native sorts of chickens are expecting a significant job in provincial economies. Native fowl execution can be improved by a modification in cultivating, taking care of, and better prosperity spread. However, selection and crossbreeding, or both can be used for genetic improvement. Aside from conventional breeding, molecular breeding techniques (Microsatellite markers and SNPs), functional genomics, gene silencing, and genome editing techniques can use from improving the quality traits in chickens. Microsatellite markers, SSCP, and sequencing techniques can use for the distinguishing proof of polymorphism on genes and assessment of the impact of polymorphism on growth traits in chickens. The molecular genetics have given a few useful assets to exploit the extraordinary abundance of polymorphism at the DNA level, for example, DNA based genetic markers. The advantage of mtDNA is that it is present in higher copy numbers within the cells and thusly bound to be recovered even from highly degraded specimens [11]. For species identification, forensic studies, anthropological and evolutionary research, the mitochondrial genome SNPs are assuming a significant role [69]. In 2016, single nucleotide polymorphism was observed and reported eleven SNPs in the D-loop region of chicken mitochondrial DNA [70].

To set up a phylogenetic relationship, a few investigations were made between domestic fowl, different red jungle fowl subspecies, and other jungle fowls. In 1994, an investigation was proposed the G.g. gallus is a major or sole contributor for a single domestication event and further three species of RJF mtDNA D-loop region sequences were taken for phylogenetic analysis and proved G.g. gallus is the real matriarchic origin of all the domestic poultry [5,11]. In 2009, two ribosomal genes (12S rRNA and 16S rRNA) were used to study the genetic divergence between Indian Red Jungle Fowl (G.g. murghi), Gallus gallus subspecies (G.g. spadicus, G.g. gallus and G.g. bankiva) including G.g. domesticus (domestic fowl) and three other Gallus species (G. varius, G. lafayetei, G. sonneratii) and found that, compared to other jungle fowls, G.g. murghi was more close to Gallus gallus subspecies, however between Indian RJF and other RJF subspecies, the divergence was very low [71]. Furthermore, found no noticeable contrasts among G.g. spadicus and G.g. gallus yet G.g. bankiva showed contrasts with both the genetic and phylogenetic relationships among these species [72]. Recent studies confirmed multiple domestications of Indian and other domestic chickens and suggested the evidence for the domestication of Indian birds from G.g. spadiceus, G.g. gallus, and G.g. murghi [73–75]. These studies avoided the two different subspecies of red jungle fowl i.e. G.g. murghi and G.g. jabouillei, however, provided a framework for genetic studies in wild jungle fowls, native, and domestic chicken breeds. Considering solid confirmations of domestication of chicken in Indus valley, it might be fascinating to contemplate the genetic relatedness between Indian Red Jungle Fowl and other Red Jungle Fowl subspecies including G.g. domesticus and other jungle fowls [76].

Indian RJFs are widely distributed across 51 x 10^5^ km^2^ in 21 states of India [77] (Fig. 1). We were more interested to know the phylogenetic relationship of Indian RJF with reference RJF and Asian native breeds along with other Indian native breeds and also understanding the contribution of Indian RJF, G. g. murghi to the domestication event. Hence, in the present study, an endeavor has been made with seven Indian native breeds (Aseel, Ghagus, Nicobari brown, Kadaknath, Tellicherry, Haringhata black, and Indian Red Jungle fowl) along with twenty-two Asian native breeds to study the nucleotide sequence variation in complete mtDNA and the most variable region of mtDNA D-loop region to build up the phylogenetic relationship among the Indian breeds (Table 1; Table 2). Starting at now, no sequence data is accessible from these seven Indian native breeds which, probably, was the contributor of one of the earliest known chicken domestication events, for example in the Mohanjo-Daro Indus valley.

Over various avian species, they incorporate Struthioniformes, Falconiformes, and Sphenisciformes other than Galliform, these conserved characteristics have been portrayed [78,79]. Even though the Indian indigenous chicken D-loop area sequence has the closeness of cytosines and guanines strings in proximately to each other makes the advancement of a consistent hairpin structure possible [80]. In all neighborhood chicken of India, the conserved sequence motifs of TACAT and TATAT were found. Such subjects are depicted as termination-associated sequences segments (TASs) drew in with the finish of mtDNA synthesis [81]. The both Galliformes and mammals have the TASs, it may propose a strong auxiliary capacity of the D-loop area of the two genera, while the nonattendance of variety in TASs among the Galliformes may be a direct result of the particular utilitarian constraints. The 8 haplotypes recognized from the Indian indigenous chicken and the phylogenetic examination spoke to the formative associations.

A large portion of the investigations on the domestic chicken origins was focused around the D-Loop; there were additionally a few examinations concentrating on the mitochondrial genomics to analyze the domestic chicken origins [82]. Kauai feral chicken’s mitochondrial phylogenies (D-loop and whole mtDNA) revealed two different clades within their samples [83]. Maybe the purpose behind the thing that difference was that the samples for the examination were different between whole mtDNA phylogeny and mtDNA D-loop phylogeny. We found that the mitochondrial phylogenies (D-loop and whole Mt genome) uncovered a few clades, and their examination re-evaluated the worldwide mtDNA profiles of chickens and encouraged our comprehension about the settlement in India.

From the mtDNA examination, we saw that Indian RJF is the origin of reference RJF as well as Indian breeds (i.e. Aseel, Ghagus, Nicobari brown, Kadaknath, Haringhata black and Tellicherry). The reference RJF is closely related with six Indian native breeds imparted to Indian reference RJF. This is likewise true for Indian chicken that has originated by independent domestication from Indian RJF as well as could be expected from other RJF species. Curiously, sharing of various Indian breeds with Indian RJF explain that the current day Asiatic chicken may have originated from various ancestors by numerous domestication events and such multi-origin breeds could still be seen in a single geographical location. This is reliable with the current day perceptions of native Japanese chickens and they are originated from various regions [84]. However, all the breeds aside from Indian Red Jungle Fowl form a solitary group suggesting a common ancestor long back in history for these birds including jungle fowls and domestic birds. The separation of Indian Red Jungle Fowl from the main group of breeds shows the possibility of a speciation event. Our examination uncovered that the Indian breeds are relatively pure with uncommon hybridization with reference RJF. In the current investigation of Indian breeds, we did not come across recognizable hybridization at least in the recent past, as shown by a clear separation of Indian RJF clades from reference RJF in mtDNA based phylogeny. All these results indicate the genetic integrity of the Indian RJF.

In high-altitude birds, hypoxia is unavoidable environmental stress, in natural selection to increase the efficiency for adapting to hypoxia every step in aerobic respiration must have experienced [85–87]. Mitochondria play an important role in aerobic respiration through oxidative phosphorylation, because of the majority of produced cell ATP consumed for cellular oxygen uptake [29,30]. To determine the role of mitochondrial genes in high-altitude adaptation, 6 high-altitude Phasianidae birds, and 16 low-altitude relatives mitochondrial genomes were analyzed and found four lineages for this high-altitude habitat [31]. Their results strongly suggests that the adaptive evolution of mitochondrial genes i.e. ND2, ND4, and ATP6 played a critical role during the independent acclimatization to high altitude by galliform birds. In Tibet, a chicken breed found a missense mutation in the MT-ND5 subunit of the NADH dehydrogenase gene for high-altitude adaptation [32]. To explore the regulatory mechanisms for hypoxia adaptability, the ATP-6 gene was sequenced from 28 Tibetan chickens and 29 Chinese domestic chickens and detected six SNPs [35]. In high-altitude adaption, cytochrome c oxidase (COX) was the key mitochondrial gene and play an important role in oxidative phosphorylation regulation and oxygen sensing transfer. For identifying the COX gene SNP, Tibet Chicken and four lowland chicken breeds (Dongxiang Chicken, Silky Chicken, Hubbard ISA White broiler, and Leghorn layer) were used and 13 haplotypes were defined for the 14 SNPs and concluded that the significant difference in MT-CO3 gene mutation might have a relationship with the high-altitude adaptation [33]. MT-CO3 gene was sequenced by using 125 Tibetan chickens and 144 Chinese domestic chickens, identified eight SNPs, and was defined into nine haplotypes, and they found positive and negative haplotype association with high-altitude adaptation [34]. However, MT-COI gene was sequenced from 29 Tibetan chicken, and 30 Chinese domestic chickens and 9 SNPs were detected [35]. In our study, ND3, ND4, ND5, ATP6, COX I, and COX-II showed high-altitude lineages between Nicobari brown and Reference RJF birds and may help evolve to adapt in Nicobari Islands environments (Fig. 5; Fig. 6; Fig. 7; Fig. 8).

For a different scope of human diseases, mtDNA mutations contribute to enclosing both tissue-specific and multiple system disorders [88,89]. The spindle-associated chromosomal exchange didn’t show antagonistic consequences for fertilization on subsequent embryo/ fetal development in the rhesus monkey [90]. Subsequently, this method may speak to another dependable restorative way to deal with the transmission of mtDNA mutations in influenced families. Mitochondrial heterogeneity is the presence of at least two kinds of mitochondrial (mt) DNA in the same individual/tissue/cell and it is firmly related to animal health and disease. In mtDNA, ND2 is a protein-coding gene and it is partaking in the mitochondrial respiratory chain and oxidative phosphorylation. In cloned sequencing of the ND2 region, numerous potential heteroplasmic locales were recognized, which were possibly reflected bountiful heteroplasmy in the chicken mitochondrial genome [91]. These outcomes give a significant reference for further research on heteroplasmy in chicken mitochondria. As of late, an examination was utilized complete mtDNA from tuberculosis patient’s blood samples and explored the conceivable mtDNA variations [92]. Twenty-eight non-synonymous variants were found and most of the variations lie in the D-loop of the non-protein-coding region of the mitochondrial DNA. Runting and stunting syndrome (RSS) generally happen early in life and causes low body weight, and prompts extensive economic losses in the commercial broiler industry [93]. In sex-linked dwarf (SLD) chickens, the RSS is related to mitochondria dysfunction, and mutations in the TWNK gene are one reason for mtDNA exhaustion [94,95]. We recommend that mutations in the mitochondrial genome should be validated further to comprehend their relationship with animal diseases.

## 5. Conclusions

In summary, this current study revealed insight into the phylogenetic relationship of Indian native chicken breeds using entire mtDNA and D-loop sequences. Eight haplotypes of the native chicken population indicated a relatively rich genetic pool. The neighbor-joining tree using genetic distance showed that the South East Asian RJF is distant to the Indian RJF but genetically close to the Indian breeds. Also, Indian Aseel breed was more closely related to the reference RJF than the Indian RJF. The grouping of Indian RJF separated from Indian native chickens and the presence of subcontinent explicit haplogroups gives an additional proof for an independent domestication event of chicken in the subcontinent. The phylogenetic analysis showed the genetic relationship within the Indian breeds and molecular information on genetic diversity revealed may be useful in developing genetic improvement and conservation strategies to better utilize precious genetic reserve. For high-altitude hypoxic adaptation, it is important to improve the efficiency of oxygen usage instead of enhancing oxygen uptake and transport. Thus, in natural selection the mitochondrial genome encoded 12 essential structural genes (6 NADH dehydrogenase genes, cytochrome b subunit, 3 cytochrome c oxidase, and ATP synthase subunit), must have mutated during adaptation to high-altitude hypoxic conditions.

## Author’s Contribution

KM analysed the data and prepared the draft manuscript. TKB developed the idea, planned the research work, carried out the wet lab experiment and edited the draft. RNC analysed the data and prepared the tables and graphs. UR provided Aseel sample, SH provided Ghagus, Nicobari (Bkack and Brown) and Kadaknath samples, MRR collected the samples of chicken breeds.

## Acknowledgements

The authors thankfully acknowledge the WBUAFS, West Bengal and CSKHPKVV, Palampur for providing blood samples to carry out the research work. Corresponding author also convey thanks to the Department of Science & Technology (SERB) (Project No. CRG/2018/002246), Government of India for providing financial support to carry out the research work.

## List of abbreviations

NGS: Next generation sequencing
OXPHOS: oxidative phosphorylation
mtDNA: Mitochondrial DNA
NAD1: NADH dehydrogenase subunit 1
NAD2: NADH dehydrogenase subunit 2
NAD3: NADH dehydrogenase subunit 3
NAD4: NADH dehydrogenase subunit 4
NAD5: NADH dehydrogenase subunit 5
NAD6: NADH dehydrogenase subunit 6
ATP6: mitochondrial encoded ATP synthase membrane subunit 6
COX1: cytochrome c oxidase subunit I
COX2: cytochrome c oxidase subunit II
COX3: cytochrome c oxidase subunit III
CYTB: Cytochrome b
rRNA: Ribosomal RNA
SNP: Single-nucleotide polymorphism
INDELs: Insertions and deletions
SEA: South East Asia
RJF: Red Jungle Fowl
ICAR-NBAGR: Indian council for agricultural research-National bureau of animal genetic resources
RSS: Runting and stunting syndrome
SLD: sex-linked dwarf
TWNK: Twinkle mtDNA helicase

## Supporting information

**S1 Fig.** PCR products loaded on 1% agarose gel. Samples A1-A11-Aseel; A13-A23-Ghagus; B1-B11-Nicobari brown; B13-B23-Nicobari black; C1-C11-Tellichery; C13-C23-Kadaknath; D1-D11-Haringhata; D13-D23-Red Jungle Fowl; A12, B12, C12, D12 are ladder.

**S1 Table.** Single nucleotide variants (SNVs) called in whole mtDNA sequencing analysis of samples from eight Indian native chicken breeds. The results from sequencing 500ng total mtDNA extracted from whole blood.

**S2 Table.** Organization of the mitochondrial genome in seven Indian native chicken breeds.

